# Interaction between TREM2-Macrophages and *Cutibacterium acnes* Drives Altered Lipid Metabolism in Chronic Apical Periodontitis

**DOI:** 10.64898/2026.04.19.719526

**Authors:** Dawool Han, Nam Hee Kim, Seung-Yong Han, Hyeeun Song, Sue Bean Cho, So Young Cha, Yechan Baek, Kyeongyeon Kim, Sungryung Kim, Hyounmin Kim, Jin-Young Park, Jung-Seok Lee, Sung Jae Shin, Su-Hwa Lee, Eunae Sandra Cho, Hyun Sil Kim, Dohyun Kim, Jong In Yook

**Author notes:** These authors contributed equally to this work (Dawool Han, Nam Hee Kim). Correspondence (J.I.Y) or (H.S.K) or (D.H.K).

## Abstract

Chronic periapical periodontitis (CAP), highly prevalent worldwide, has long been regarded as non-specific inflammation. Lipophilic *Cutibacerium acnes* (CA) persistence in host macrophages has emerged as the pathologic background of sarcoidosis and acne vulgaris. Here we report that intracellular persistence of CA in TREM2-macrophages plays a pathologic role in CAP. We observed persistent CA in macrophages in most CAP samples. Our CA clinical isolates persist in the cytosolic space of macrophages, retarding phagolysosomal degradation accompanied by NLRP3-dependent inflammatory response. Subcutaneous injection of those isolates *in vivo* recapitulates subcutaneous aggregation of CA-laden macrophages. By single cell RNA sequencing analysis of defined CAP samples, we found that CA in TREM2-macrophages drives exuberant lipid droplets formation, indicating that immune cells are potential lipid provider in CAP. Our observations elucidate the mechanistic link whereby TREM2-macrophages and altered lipid metabolism provide a lipid-rich niche for CA, contributing to the pathophysiology of CAP.

---

Chronic apical periodontitis (CAP) around a tooth’s root apex is a global health issue impacting the overall health and socioeconomic status of affected individuals^1^. CAP typically develops from the exposure of the dental pulp to oral microbiota via dental caries or accidental trauma. The spread of microorganisms through dental pulp into the periapical region of the affected tooth results in chronic inflammation, bone resorption, and destruction of surrounding tissues, frequently resulting in tooth loss^2^. Although routine non-surgical root canal treatment has a high success rate, the failures followed by treatment-refractory CAP running as high as 20%^3^. Clinically, CAP is characterized by long-standing (months to years) intrabony inflammation and is refractory to treatment of antibiotics. Histopathologically, CAP has long been regarded as non-specific inflammation because causal microorganisms remain unidentified^4–6^.

Commensal *Cutibacterium Acnes* (CA, formerly *Propionibacterium acnes*) in skin and oral mucosa is anaerobic, gram-positive, and a well-known lipophilic bacterium. Although CA is low-virulent, it is closely linked to many human intractable diseases, including skin acne vulgaris, prostate cancer, chronic ocular and prosthetic inflammations^7–9^. Interestingly, intracellular CA persistence has emerged as having a potentially causative role in sarcoidosis^9,10^, a disease characterized by granuloma (cluster of activated macrophages). Histopathologic and immunohistochemical analysis reveals latent infection of cell-wall-deficient CA in macrophages (M□) of sarcoidosis lesions, similar to *Mycobacterium tuberculosis* (*Mtb*)^9,11^. Intratracheal administration of CA isolated from sarcoidosis lymph nodes successfully generated pulmonary granuloma in an innate immunity-dependent manner^12,13^. While various intracellular pathogens, especially in innate immune cells, adopt specific interactions with a host and/or a microenvironment^14,15^, host-microbiome interaction between M□ and CA followed by micro-environmental change is not fully understood.

TREM2 (triggering receptor expressed on myeloid cells 2) is a surface receptor wildly expressed on monocyte-lineage immune cells. The importance of TREM2 as a key sensor of lipid homeostasis is well documented in obesity, neurodegenerative, and other inflammatory diseases mediating diverse functions in phagocytosis, inflammatory signaling, survival, and proliferation^16–19^. In acne vulgaris, a skin lesion affecting more than 45 million people, the host-microbe interaction between CA and TREM2^high^-macrophages (TREM2-M□) induces altered lipid metabolism and a proinflammatory response^20^, indicating that CA persistence in TREM2-M□ plays an important role in inflammatory response via alteration of lipid metabolism.

Previously, we reported the persistence of CA in M□ of peri-implantitis, a treatment refractory dental inflammation which has emerged due to the popularity of dental implants^21^. Based on our previous observations, we hypothesize that intracellular CA persistence occurs in other chronic inflammatory lesions. We found intracellular CA persistence in M□ among the majority of CAP clinical samples. *In vitro*, our clinical isolates persist in the cytosolic space of M□, retarding or escaping phagolysosomal degradation concomitant with a strong acute inflammatory response in a NLRP3-dependent manner. Subcutaneous injection of our isolates *in vivo* generates granuloma-like aggregation of CA-laden M□, indicating that innate immune cells are major hosts for CA. Single cell RNA sequencing (scRNA-seq) analysis of CAP samples identified TREM2-M□ playing an important role in lipid metabolism of M□. Notably, intracellular CA was co-localized with TREM2 in human PBMC-derived M□ (hPBMC-M□) and in clinical samples of CAP, indicating that CA persistence can inhibit TREM2 function^22^. Indeed, CA-infected hPBMC-M□ or TREM2 knockout (KO) in THP-1 cells induced formation of abundant lipid droplets (LD), indicating that immune cells can be a lipid provider for niche of CA in CAP. Consequently, we investigated altered LD accumulation regulated by host-microbiome interactions between TREM2-M□ and CA underlying the CAP pathophysiology.

## Results

### Identification and isolation of intracellular CA from CAP lesions

Building on our previous observation of CA in extensive peri-implantitis lesions^21^, we chose more prevalent CAP clinical samples. Considering the wide range of oral microbiomes, anatomic complexity of periapical tissues, and variable etiology of periapical inflammation, in this study we define CAP as occurring in endodontic microsurgery cases (known as apical surgery) following failure of conventional root canal treatment, also known as post-treatment apical periodontitis^2,23^. We initially used routine immunohistochemical (IHC) staining with PAB antibody, a monoclonal antibody specific to lipoteichoic acid of C. acnes^24^. Among 7 cases of retrospective formalin-fixed paraffin-embedded (FFPE) samples of CAP (Supple. Table 1), we found intracellular CA to a varying extent in all cases, especially in the cytosolic space of inflammatory cells (Fig. 1a and Extended Data Fig. 1a). In many cases, we found several dozen CA in the cytosolic space of immune cells. Multiplex immunohistochemistry (mIHC) was employed to further verify CA localization in a single section, using anti-CD68 for endo-lysosomes, anti-CD14 for surface markers of M□, and PAB antibody for CA. Indeed, CA in CD68 and/or CD14-positive M□ was detected in all clinical FFPE samples (Fig. 1b and Extended Data Fig. 2a), while the population and distribution of CA were largely variable. Though most CA were co-localized with CD68^+^ lysosomes, many intracellular CA in CAP samples were not in lysosomal space^25^. We next generated polyclonal antibody (anti-CA) using intact CA isolate (Extended Data Fig. 1b) to further evaluate the specificity of intracellular CA in clinical samples. We performed an immunofluorescence (IF) study with anti-CA and PAB antibodies using M□-enriched CAP clinical samples using magnetic bead sorting (MACS) with CD14 antibody, detecting intracellular CA in CD68^+^ M□ in 4 out of 5 cases by means of anti-CA and anti-PAB antibodies (Fig. 1c). These observations indicate that intracellular CA may play an important role in the pathogenesis of CAP.

**Fig 1.**
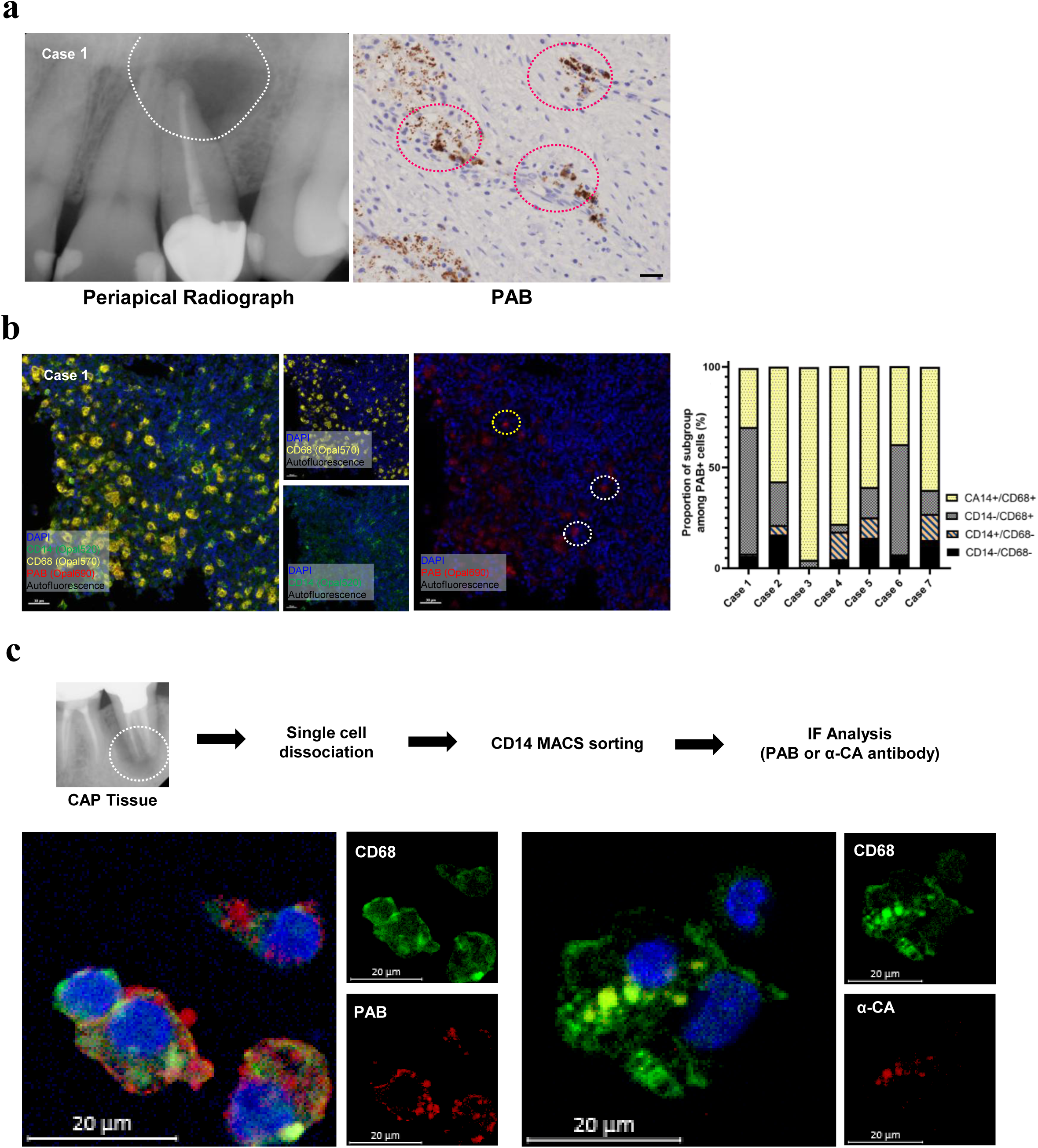
Intracellular CA persistence in M□ in CAP lesions. **a. Detection of CA in CAP lesions.** Representative periapical radiograph showing a radiolucent lesion (white dotted line) at the apex of tooth #22 (left panel). Representative IHC staining using a PAB antibody revealed the presence of CA as small punctate signals within the cytoplasm of inflammatory cells (red dotted circles). (Scale bar, 30 μm) **b. Multiplex immunohistochemical (mIHC) analysis of CA localization in M**□. mIHC was performed using antibodies against CD14 (surface marker), CD68 (endolysosomal marker), and CA (PAB) in retrospective FFPE tissues from seven cases of CAP. Representative images show CA signals (red) localized within CD14□ and/or CD68□ cells (white dotted circles), as well as signals detected outside CD14□/CD68□ cells (yellow dotted circle). The proportion of CA in M□ (CD14+/CD68+, CD14-/CD68+, CD14+/CD68-) and out of M□ (CD14-/CD68-) was automatically counted as described in Methods (right panel). Note a few proportion of CA is localized in outside of M□ (black). (Scale bars, 30 μm) **c. Validation of intracellular CA in M**□ **using a strain-specific antibody.** Schematic illustration of the experimental workflow for enrichment of CD14□ monocyte/macrophage populations from CAP tissues, followed by immunofluorescence (IF) staining (upper panel). CAP samples were dissociated into single cells, and CD14□ cells were isolated by MACS prior to IF analysis. Representative confocal images show CA signals detected using both PAB and strain-derived polyclonal anti-CA antibodies within CD68□ cells. (Scale bars, 20 μm)

To elucidate the role of CA in dental inflammation, we obtained two clinical CA isolates (D1 and D2) using MACS with CD14 antibody from peri-implantitis and CAP lesions, respectively (Supple. Table 2), followed by 16S rRNA sequencing to identify those isolates as CA. We also obtained C1 isolate originated from sarcoidosis lesion (kindly provided by Professor Eishi). Phylogenetic analysis using high-resolution single locus sequence typing (SLST)^26^ revealed that our D1 and D2 isolates were characterized by clusters K2 (type II) and L3 (type III), respectively (Extended Data Fig. 3a). Interestingly, those K and L types are mainly found in the oral cavity, while C1 isolate was identified in cluster A5 (type 1A) the subtype found mainly in skin^26,27^. An antimicrobial susceptibility test revealed that our isolates are susceptible to penicillins and 2^nd^ generation cephalosporins as well as resistant to metronidazole and chloramphenicol, consistent with previous observations related to the treatment of acne vulgaris^28^ (Extended Data Fig. 3b).

### Persistent localization of CA in M**□** escaping phagolysomal maturation

M□ constitute the primary innate immune cells with extensive antibacterial phagocytic activity. Although the pathologic roles of intracellular CA in M□ in many inflammatory lesions, such as acne vulgaris and sarcoidosis, are well-known, the host inflammatory response to and intracellular fate of CA in M□ have not been fully elucidated^25,29^. Prior to conducting CA experiments in an *in vitro* model, we validated the active phagolysosomal capacity of RAW264.7, THP-1 cells and hPBMC-M□ using GFP-expressing *E. coli*, finding rapid phagolysosomal maturation (within an hour) of engulfed bacteria (Extended Data Video 1, 2 & 3). Previously, THP-1 cells infected with prostate CA isolate at an MOI of 25 and RAW264.7 cells infected with CA isolate up to MOI of 100 revealed viabilities of both host cells and intracellular CA for 3 days^25,29^. To examine the host cell cytotoxicity in terms of intracellular uptake and persistence of CA, we treated our isolate into M□ up to MOI of 100, then measured host cell viability and intracellular persistence of CA. As in previous observations^25,29^, hPMMC-M□ and RAW264.7 cells were viable and tolerable treated up to MOI of 100 (Extended Data Fig. 4a), suggesting the low-virulent character of CA in M□. We next treated CA isolates into the hPBMC-M□ and examined intracellular fate using IF staining (Fig. 2a). Interestingly, we detected the intracellular presence of CA, mainly in the juxta-lysosomal space of hPBMC-M□ for 5 days and RAW264.7 cells for a 10-day period (Fig. 2b and Extended Data Fig. 4b). While several CA were co-localized with lysosomes, most persisted in the cytoplasm of M□ regardless of bacterial viability (Extended Data Fig. 4c), indicating that CA persists in the cytoplasmic space for a long period, escaping from phagolysosomal maturation. Because intracellular CA are viable in RAW264.7 and HeLa cells^29^, we treated our isolates to hPBMC-M□ for 2 days and plated the cytosolic lysate of macrophages for colony formation assay. Indeed, our isolates were also viable for 2 days after recovery from hPBMC-M□ and able to replicate on anaerobic agar plates whereas heat-killed CA (HK-CA) isolate was not (Extended Data Fig. 4d).

**Fig. 2.**
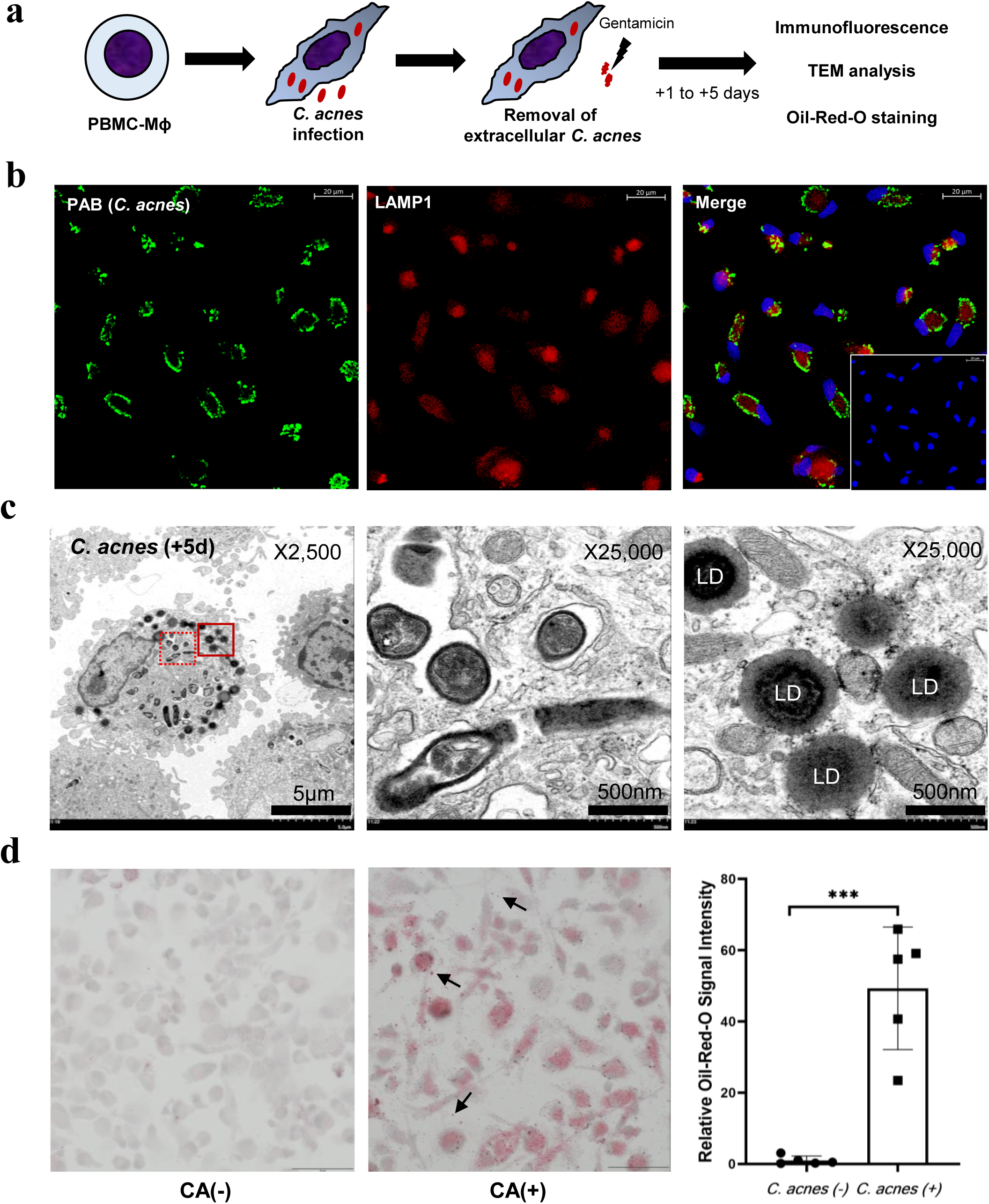
Delayed phagolysosomal maturation of intracellular CA induces abundant LD formation. **a. Experimental design for analysis of intracellular fate and host responses to CA.** hPBMC-M□ were infected with CA, and intracellular localization, persistence, and host responses were analyzed at indicated time points using immunofluorescence, transmission electron microscopy, and lipid staining assays. **b.** Intracellular localization of CA in M**□** relative to lysosomal compartments. Representative immunofluorescence images hPBMC-M□ infected with CA. CA (red) was detected within M□ and showed partial co-localization with lysosomal marker CD68 (green). While a subset of bacteria overlapped with CD68□ compartments, a substantial fraction of intracellular CA was observed outside lysosomal regions, indicating localization in the cytoplasmic or juxta-lysosomal space. (Scale bars, 20 μm) **c. Ultrastructural analysis of intracellular CA persistence in M**□. TEM images of hPBMC-M□ infected with CA (MOI 40) and examined at 5 days post-infection. Intracellular bacteria observed within membrane-bound compartments exhibited preserved structural integrity, including identifiable cell wall and electron-dense cytoplasmic features, suggesting limited degradation (dotted rectangle, middle). In addition, infected cells showed an increase in lipid droplets (LD) structures in the peripheral cytoplasm (lined rectangle, right). (Scale bars, 5 μm (left) and 500 nm (middle and right)) **d. Increased lipid accumulation in M□ following CA infection.** Representative images of Oil Red O staining in hPBMC-M□ infected with CA (MOI 20) and analyzed at 48 hours post-infection. CA-infected cells (middle panel) exhibited increased intracellular lipid droplet formation compared to uninfected controls (left panel). Note multiple LD in extracellular space of M□ (arrows). For quantification, Oil Red O–positive signals were extracted and measured based on red channel intensity using ImageJ software. Data are presented as relative lipid accumulation. (Scale bars, 50 μm)

### TEM analysis reveals the variable status of CA and abundant LD formation in M**□**

To evaluate the ultrastructure of intracellular CA, we next performed TEM analysis of CA-infected M□. Similar to HeLa cell observations^29^, TEM analysis with hPBMC-M□ at 3 days and 5 days post-infection (MOI 40) showed variable structures containing digested and undigested CA (Fig. 2c and Extended Data Fig. 4e). We found CA to be localized in membrane-bound compartments as well as in the cytosolic space of hPBMC-M□. Sporadically, CA was identified in multivesicular bodies in variable digested states. In contrast to HeLa cells^29^, persistent CA in hPBMC-M□ harbors many LDs in proximity to intracellular CA (Fig. 2c). Abundant LDs were found 3 days post-infection in viable as well as in HK-CA (Extended Data Fig. 4e), suggesting that CA persistence induces LD formation by M□. To further evaluate the role of CA in lipid metabolism, we treated CA isolate into hPBMC-M□ and examined LD formation by means of Oil Red O staining. Indeed, CA persistence produces abundant LDs in the cytosolic space as well as in extracellular space of hPBMC-M□ *in vitro* (Fig. 2d), pointing to immune cells as a potential provider of lipid in CA-linked lesions.

### Inflammatory response to CA persistence in M**□**

While acne vulgaris constitutes an important health issue driven by CA, a large number of biological response studies have focused on epithelial cells or sebocytes^20,30,31^, studies on the inflammatory response of immune cells being notably fewer. To compare transcriptional responses of our isolates in M□, we performed a preliminary bulk RNA sequencing study with C1, D1, and D2 isolates using RAW264.7 cells and DEG analysis. We found that those isolates produced similarly strong inflammatory responses with Il1α, Il1β, Il6, Ccl2, and Il23 (Extended Data Fig. 5a), suggesting few differences among those CA isolates. To examine the acute inflammatory response, we next treated our CA D1 isolate in hPBMC-M□ for 24 h and subjected it to bulk RNA sequencing (RNA-seq). DEG analysis of CA-infected M□ revealed strong inflammatory responses to IL1α, IL1β, IL23α, IL6, MMPs and chemokines, suggesting an acute M1-like inflammatory response (Fig. 3a). We next performed KEGG pathway enrichment and gene ontology (GO) analysis to identify enriched biological functions and networks from RNA-seq. KEGG and GO enrichment analysis revealed that the up-regulated DEG were involved in diverse inflammatory pathways, such as cytokine-receptor interaction, lipid-atherosclerosis, tuberculosis and TNF-α signaling (Extended Data Fig. 5b). To validate transcriptional response, we next performed qRT-PCR analysis of selected genes. Transcriptional abundance of those acute inflammatory genes was highly increased by CA treatment in hPBMC-M□ for 5 days post-infection (Fig. 3b). Measuring the secreted IL1β level, we found it significantly increased by CA in hPBMC-M□ (Fig. 3c). Because the NLRP3-inflammasome is a critical component of the innate immune system activating IL-1β and IL-18^32^, we next generated NLRP3 knockout (NLRP3-KO) THP-1 cells to determine the role of the NLRP3-inflammasome in CA-driven IL1β secretion. We found this secretion to be largely dependent on NLRP3 in THP-1 cells (Fig. 3d and Extended Data Fig. 5c). A multiplexed cytokine assay of the hPBMC-M□ culture medium revealed that CA isolates significantly increased acute inflammatory cytokines such as IL6, GM-CSF, CCL8 (known as MCP2), CCL3 (MIP1α), and CCL15 (MIP1δ) (Fig. 3e and Extended Data Fig. 5d). These results indicate that CA persistence strongly induced NLRP3-dependent acute inflammation in M□.

**Fig. 3.**
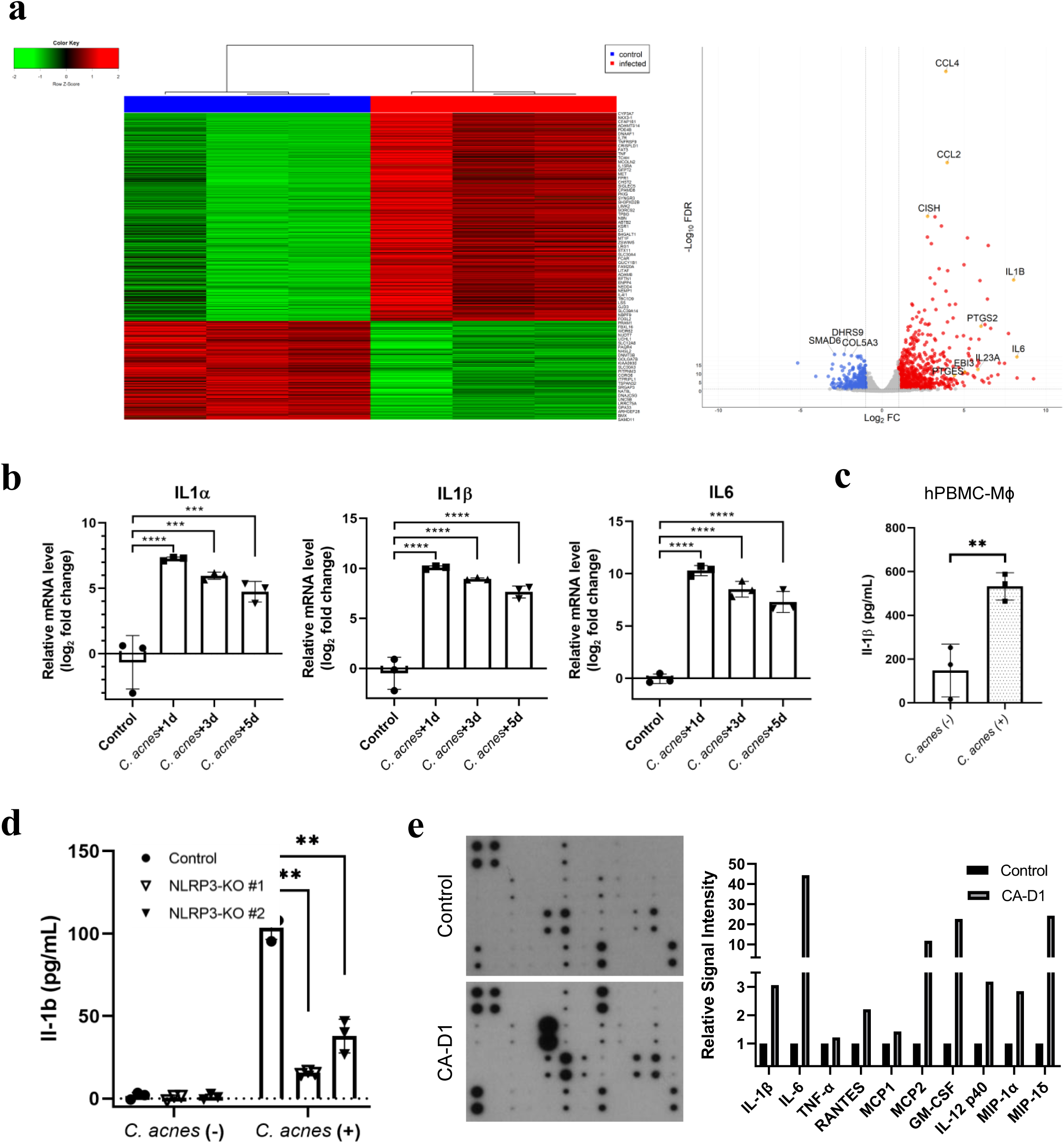
Acute inflammatory response of hPBMC-M□ by CA. **a. Transcriptomic profiling of M**□ **following CA infection.** hPBMC-M□ were infected with CA (CA-D1, MOI 20) for 24 hours, followed by bulk RNA sequencing analysis. Heatmap showing differentially expressed genes (DEGs) demonstrates distinct clustering between control and infected samples (left panel). Volcano plot illustrates significantly upregulated and downregulated genes, with notable induction of pro-inflammatory cytokines and mediators, including IL1α, IL1β, IL6, IL23α, and chemokines such as CCL2 and CCL4, consistent with an acute M1-like inflammatory response (right panel). **b. Validation of inflammatory gene expression by qRT-PCR.** hPBMC-M□ were infected with CA (MOI 20), and total RNA was collected at 1, 3, and 5 days post-infection. Relative mRNA expression levels of IL1α, IL1β, and IL6 were determined by qRT-PCR (n = 3) and are presented as log□ fold change compared to control. Sustained upregulation of pro-inflammatory cytokine genes was observed up to 5 days post-infection. **c. Increased secretion of IL1**β **following CA infection.** hPBMC-M□ were infected with CA (MOI 20), and culture supernatants were collected at 24 hours post-infection. IL1β levels in the culture medium were measured by ELISA (n = 3). **d. NLRP3-dependent IL1**β **secretion in response to CA.** THP-1 cells with knockout of NLRP3 (NLRP3-KO) and corresponding control cells were treated with CA, and IL1β levels in culture supernatants were measured. CA-induced IL1β secretion was markedly reduced in NLRP3-KO cells compared to control cells (n = 3). **e. Cytokine profiling of M**□ **response to CA.** Cytokine expression in culture supernatants from hPBMC-M□ infected with CA-D1 was analyzed using a human inflammation antibody array. Representative membrane images are shown (left panels). Selected cytokines showing notable changes were quantified based on relative signal intensity using ImageJ (right).

### CA-laden M**□** aggregation in vivo

Previously, pulmonary sarcoidosis by CA was successfully generated in mouse model by repeated exposure in wild type mice^12,13^. Given the inflammatory response induced by intracellular persistence of CA, we next designed an animal experiment to determine whether the M□ are the major target of CA among the immune cells and thus a cause of the subsequent inflammatory response *in vivo* (Fig. 4a). We found that subcutaneous injection of our CA isolates generated a heavy accumulation of mononuclear immune cells with strong inflammatory response (Fig. 4b). Interestingly, we found many vacuolar spaces nearby aggregated mononuclear cells in CA injected groups compared to control, suggesting an abundant lipid accumulation *in vivo*. IHC analyses revealed that mononuclear cells mainly consisted of CD68^+^ M□ having intracellular CA (Fig. 4c). We also observed the inflammatory response of activated M□, including IL1β and TNFα (Fig. 4d). Intriguingly, HK-CA isolate also induced similar aggregation of activated M□ (Extended Data Fig. 6), suggesting the possibility that cell wall components of CA may play an important role in intracellular persistence rather than bacterial viability. Taken together, these observations indicate that M□ are the primary immune cell targets of CA-induced inflammatory responses *in vitro* and *in vivo*.

**Fig. 4.**
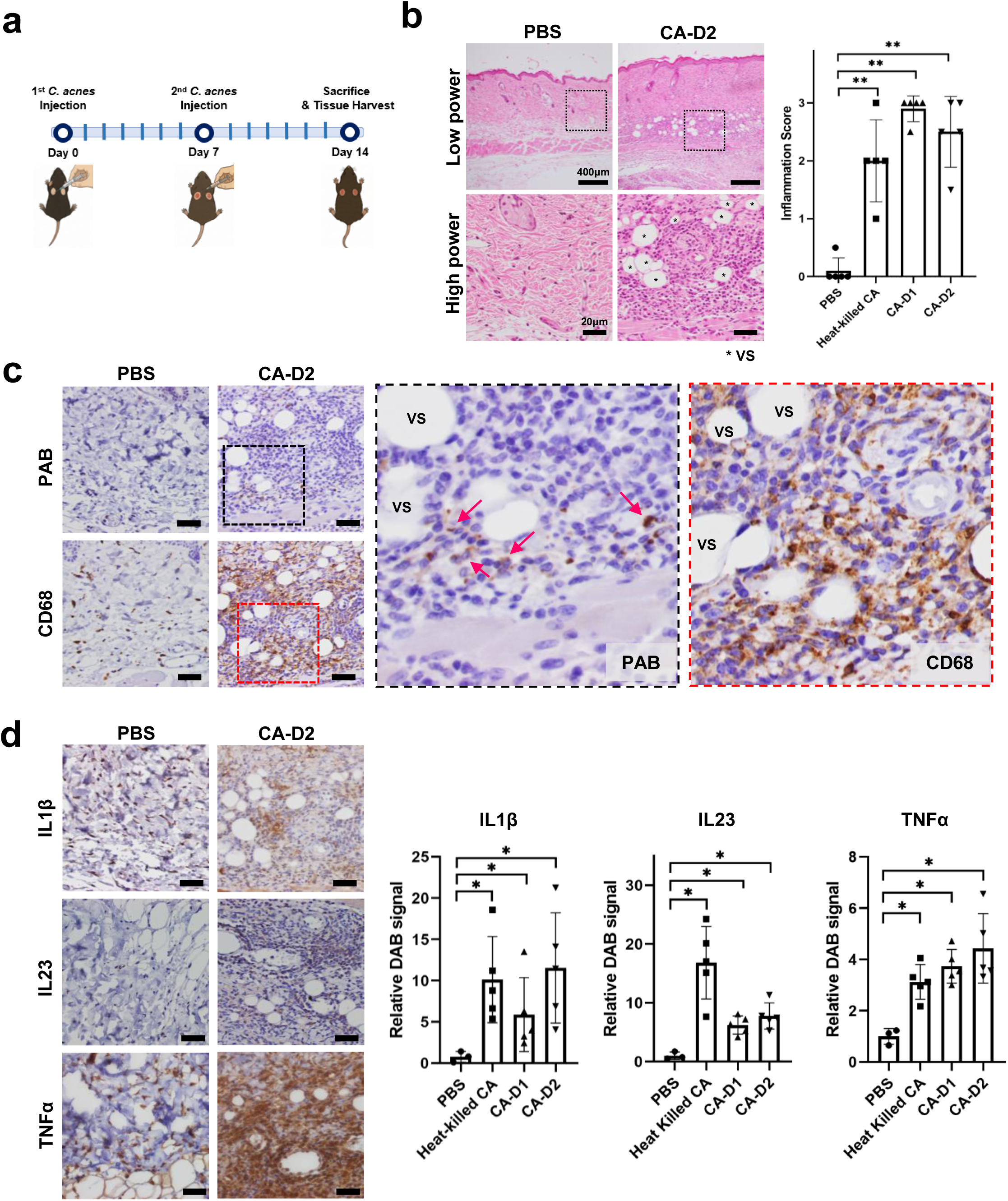
Subcutaneous injection of CA induces M□ aggregation in vivo. **a. Experimental design of the murine subcutaneous infection model.** C57BL/6 mice received two subcutaneous injections of CA at the same site 7 days apart, followed by tissue harvest 7 days after the second injection. **b. M**□**-rich inflammatory lesions induced by CA *in vivo*.** Representative hematoxylin and eosin (H&E) staining of subcutaneous tissues from mice injected with PBS or CA isolates (CA-D1, CA-D2) or HK-CA (n = 5). Representative low– and high-power images show marked accumulation of mononuclear inflammatory cells in CA-injected subcutaneous tissues compared to PBS controls. At higher magnification, vacuolated structures (*VS) were observed in areas surrounding inflammatory cell aggregates. Inflammatory responses were semi-quantitatively scored as follows: 0, no inflammation; 1, inflammatory cell infiltration; 2, aggregation of macrophages (granuloma-like structures); and 3, central necrosis. Both live and HK-CA induced comparable inflammatory responses based on this scoring system. (Scale bars, 500μm, low power and 20μm, high power) **c. Localization of CA within M**□ ***in vivo*.** Representative IHC staining of subcutaneous tissues from mice injected with PBS or CA-D2. PAB staining (top) revealed intracellular CA signals (red arrows) within inflammatory lesions, while CD68 staining (bottom) demonstrated that the majority of infiltrating cells were M□. Higher magnification images show CA signals associated with CD68□ M□-rich areas, with punctate staining observed within the cytoplasmic regions of inflammatory cells. (Scale bars, 20μm) **d. Increased inflammatory cytokine in CA-induced lesions in vivo.** IHC staining of IL1β, IL23, and TNFα in subcutaneous tissues from mice injected with PBS or CA isolates (CA-D1, CA-D2) or HK-CA. Increased cytokine expression was observed in CA-injected tissues compared to PBS controls. For quantification, DAB-positive signals were extracted and measured as relative signal intensity using ImageJ. Both live and HK-CA induced elevated expression of inflammatory cytokines *in vivo*. (Scale bars, 20 μm)

### scRNA-seq analysis revealed TREM2-M□ in CAP lesions

To elucidate dynamic cellular response in CAP that is refractory to conventional root canal treatment, we next used the BD Rhapsody microwell-based single cell analysis platform to design an scRNA-seq analysis of 5 clinical CAP samples selected using the following criteria (Fig. 5a): (1) a root canal treatment history by an endodontic specialist, (2) presence of CA-laden M□ in the lesion, (3) closed CAP lesion without oral communication, and (4) lack of trauma history, such as root fracture (Supple. Table 3). Due to the paucity of cells in normal periodontium, we used a publicly available scRNA dataset of normal cells obtained from the apical two-thirds of the tooth roots of five extracted third molars and dental pulp^33^. After excluding low-quality cells (defined as cells with >60% mitochondrial gene expression or detection of ambient hemoglobin genes) and doublets/multiplets, a total of 124,860 cells from CAP and 21,182 cells from normal periodontium and dental pulp were retained for scRNA-seq analysis. Unsupervised clustering in two-dimensional UMAP (Uniform Manifold Approximation and Projection) space identified 11 distinct clusters across 146,042 cells, corresponding to major cell types of lymphoid cells, myeloid cells, and stromal cells (Fig. 5b and Extended Data Fig. 7a). Cells from clusters of immune cells (lesional macrophages, B cells, T cells and plasma cells) were abundant in CAP samples, while the stromal cells were mainly found in healthy control (Fig. 5c and Extended Data Fig. 7b). The stromal cells included endothelial cells (characterized by the expression of genes such as SPARCL1 and TM4SF1), mural cells consisting of pericytes and smooth muscle cells (RGS5 and ACTA2), Schwann cells (S100B and MPZ) and fibroblasts (COL1A1 and COL1A2). In immune cells, we distinguished predominant plasma cells (IGHGP and IGHG1), T cells (IL32 and TRBC2), B cells (MS4A1 and BANK1), myeloid cells (LYZ and CTSB), neutrophils (CXCL8, G0S2, and S100A8), and mast cells (TPSB2, TPSAB1 and CPA3) (Extended Data Fig. 7c). Interestingly, we found HMGA1^high^ proliferating immune cells featuring B and plasma cell markers (Extended Data Fig. 8a and 8b), suggesting B-cell lineage proliferation and the existence of extrafollicular plasmablasts in CAP lesions. Further analyzing proliferating cells in CAP, we found immune cells expressing TNF family member B-cell activating factor (BAFF), a critical cytokine that selectively promotes the survival, differentiation, and maturation of memory B cells into antibody-secreting plasmablasts^34^ (Extended Data Fig. 8c). Proliferating cells are also characterized by BAFF receptor (TNFRSF13C) and B-cell maturation antigen (BCMA also known as TNFRSF17), while neutrophils and myeloid cells strongly expressed BAFF (TNFSF13B) which is also noted in acne vulgaris^20^. To validate proliferating immune cells in clinical samples, we performed IHC for Ki-67 protein, which is expressed during active cell cycle phases. In clinical CAP samples, we found varying proportions of Ki-67 positive cells in all CAP cases (Extended Data Fig. 8d). These results indicate the existence of proliferating extrafollicular plasmablasts in CAP lesions. We next focused on the myeloid cluster, given the importance of innate immune cells in CA persistence. The subclustering of the 5,271 myeloid cells identified seven populations (Fig. 5d). The three DC subclusters were defined as plasmacytoid DC (pDC of GZMB and IRF4) as an acute responder, conventional DC of cDC1 (LAMP3 and CCR7) and cDC2 (CCND2 and CLEC10A) presenting antigens to prime CD8 and CD4 T cells, respectively^35^. We also annotated lesional M□, migratory M□, and monocyte subclusters based on gene expression profiles (Fig. 5e and Extended Data Fig. 9). Regarding CAP as an intrabony inflammation, we further identified an osteoclast (OC) subset expressing the specific bone-resorbing gene SPP1 (osteopontin) and the bone matrix degradation enzyme MMP9.

**Fig. 5.**
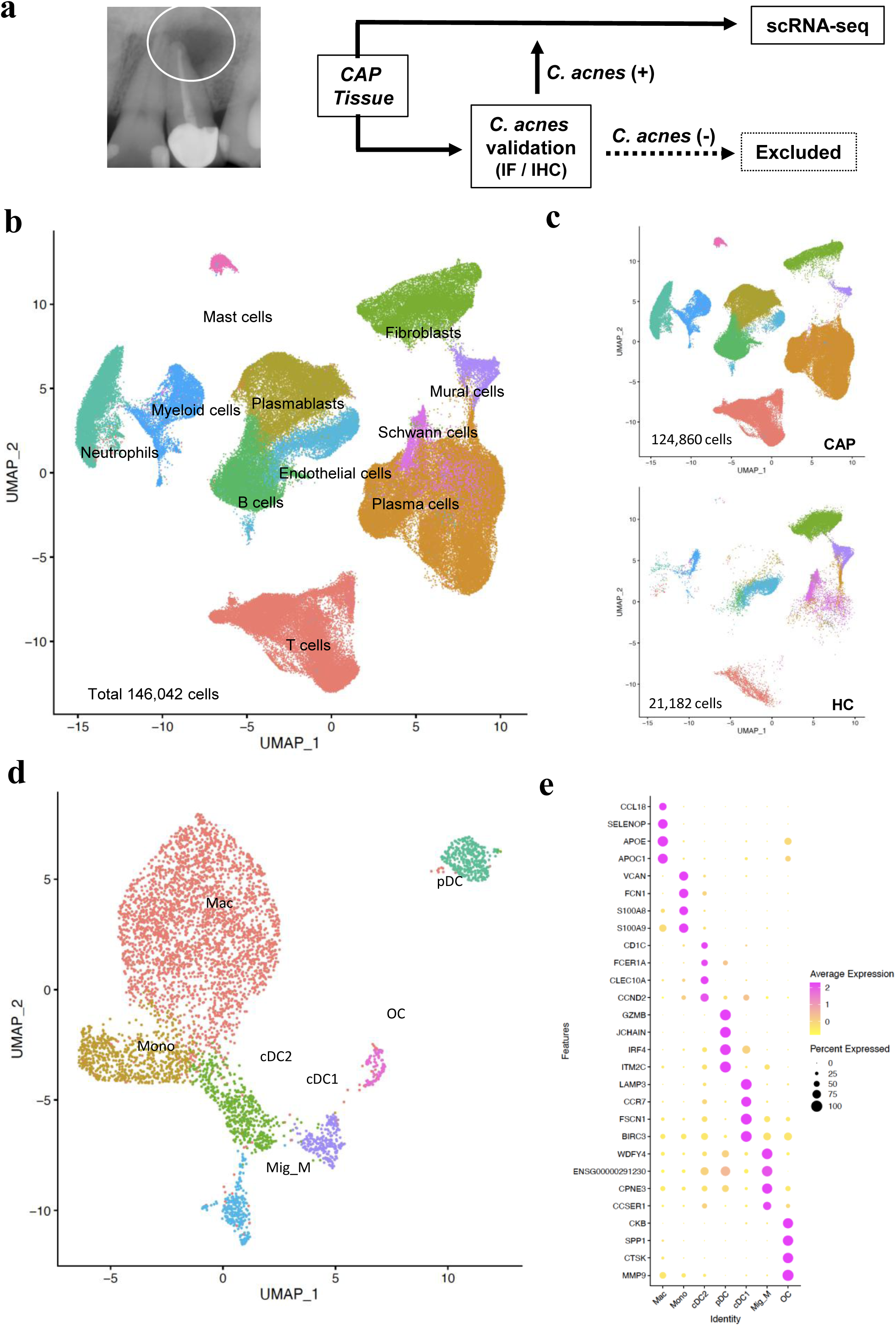
Cell types recovered from scRNA-seq analysis of CAP and healthy control. **a. Schematic illustration of scRNA-seq for CAP lesions**. Clinical samples were validated as CA using IF and IHC staining, followed by cell dissociation and scRNA-seq analysis. **b. UMAP embedding of 146,042 cells from CAP lesions and healthy control tissues, colored by cell type annotation.** Major cell populations include T cells, B cells, plasma cells, plasmablasts, myeloid cells, neutrophils, mast cells, endothelial cells, mural cells, fibroblasts, and Schwann cells. **c. UMAP plot for cell clusters from patients with CAP (upper panel) and healthy control (lower panel)**. **d. UMAP embedding of 5,271 myeloid cells from CAP lesions, highlighting major myeloid subsets including monocytes, macrophages, dendritic cells (cDC1, cDC2, pDC), migratory macrophages, and osteoclasts**. **e. Dot plot heatmap showing expression of selected marker genes for the cell of myeloid subclusters.**

### Co-localized CA in TREM2-M□ alters lipid metabolism in CAP lesions

Lipid-laden TREM2-M□s have emerged as an important subset of immune cells in Alzheimer disease, tumors, atherosclerosis, sarcoidosis and acne vulgaris^16–19^. Given the LD formation induced by CA persistence in M□, we focused on the role of TREM2-M□ and lipid metabolism in CAP. Further analysis of monocyte-M□ lineage across all myeloid clusters identified distinct macrophage subsets, including IL1β-, CCL18– and TREM2-high M□, as well as FOLR2-high tissue resident M□ (Extended Data Fig. 10a). Pseudotime trajectory analysis of myeloid subclusters revealed a continuous and directional linear trajectory from monocytes to IL1β-high macrophages, followed by transition to CCL18– and subsequently TREM2-high macrophage subsets (Fig. 6a), indicating a sequential differentiation process along a continuum of M□ states. Consistent with the acute inflammatory response observed in RNA-seq of hPBMC-M□, early inflammatory genes (IL1β, CXCL2, and CXCL8) are abundant in the IL1β-high subset (Extended Data Fig. 10b). In contrast, the CCL18 subset showed increased expression of lipid metabolism-related genes (such as APOE, APOC1, PLD3, and LIPA) and chemokines associated with CD8 T cell recruitment (CCL3, CCL4)^36^. In the TREM2-M□ subset, early inflammatory genes (including IL1β) and immediate-early transcriptional regulators (such as FOSB, JUNB, KLF4 and ATF3) were markedly reduced, whereas expression of lipid metabolism genes remained elevated. Pseudotime analysis further demonstrated a progressive increase in lipid metabolic programs, including lipogenesis and cholesterol efflux, along this sequential trajectory (Fig. 6b and Supple. Table 4). Notably, lipid metabolism peaked in the CCL18 subset and subsequently declined in the TREM2 subset along pseudotime progression (Extended Data Fig. 10c and 10d). Together, these findings support a stepwise differentiation trajectory from IL1β to CCL18 and ultimately to TREM2 macrophage states, characterized by enhanced lipid metabolism and attenuation of early inflammatory responses in the TREM2 subset.

**Figure 6.**
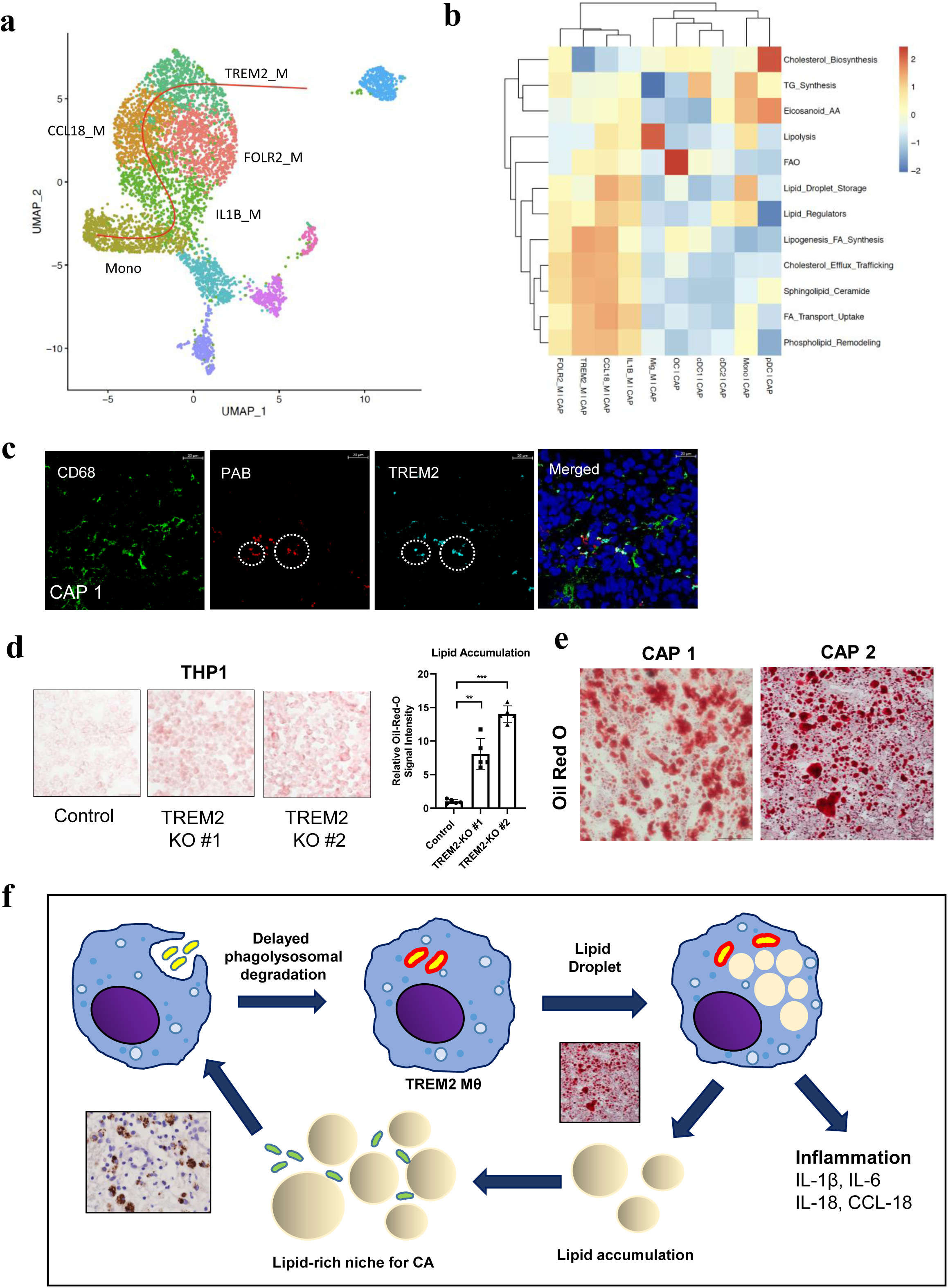
TREM2-M□ play a critical role in CA-mediated alteration of lipid metabolism. **a. Slingshot pseudotime trajectory projected onto the UMAP embedding of myeloid cells, showing a continuous and directional trajectory from monocytes to IL1β-high M□, followed by progression toward CCL18□ and TREM2□ M□ subsets**. **b. Heatmap showing gene set expression scores for lipid metabolism-related pathways across myeloid subclusters.** Scores were calculated based on the expression of genes associated with each pathway. **c. Co-localization of CA and TREM2 in macrophages detected in CAP clinical samples**. CAP samples were subjected to frozen section and IF study using anti-CD68, PAB, and anti-TREM2 antibodies. Representative IF staining in CAP samples for expression of CD68 in green, CA in red, and TREM2 in blue (CAP, n = 5). (Scale bars, 20 μm) **d. Knockout of TREM2 induced abundant LD formation in THP1 cells.** Representative images of Oil Red O staining in THP1 cells of control (left), TREM2 KO #1 (middle), TREM2 KO #2 (right). For quantification, Oil Red O-positive signals were extracted and measured based on red channel intensity using ImageJ software. Data are presented as relative lipid accumulation. (Scale bars, 50 μm) **e. Representative Oil Red O staining in CAP samples showing abundant intracellular as well as in extracellular LD**. Representative figures of Oil Red O staining using air-dried frozen sections of CAP. (Scale bars, 50 μm) **f. Graphical summary of host-microbiome interactions between CA persistence and lesional TREM2-M□ in CAP.** Our observation showed that CA inhibits TREM2 function in lesional macrophages, resulting in abundant LD formation in CAP. The lipid-rich microenviroment allows niche for lipophilic CA as well as evasion from its antibiotic susceptibility.

Given that CA-induced LD formation in PBMC-M□ and TREM2-M□ abound in scRNA-seq analyses of CAP lesions characterized by altered lipid metabolism, we next examined whether CA is responsible for TREM2-M□ observed in scRNA-seq. An IF study using CA treated PBMC-M□ revealed that intracellular CA was co-localized with TREM2 (Extended Data Fig. 10e). To further characterize the direct relationship between CA and TREM2 in lesional M□, we next conducted an IF study with CAP samples. Indeed, intracellular CA was co-localized with TREM2 in CD68^+^ M□ in CAP samples (Fig. 6c and Extended Data Fig. 10f), indicating that intracellular persistence of CA is directly related to TREM2-M□. Previous observations indicate that loss of TREM2 increased LD with loss of phagocytic activity in macrophages/microglial cells, generating foamy M□^19,37,38^. Our observations thus suggest that intracellular CA persistence may inhibit TREM2 function, resulting in altered lipid accumulation of M□. Generating TREM2-KO THP-1 cells, we found abundant LD within those cells (Fig. 10d and Extended Data Fig. 10g). These results suggest the possibility that loss of TREM2 function due to CA persistence in lesional M□ provides a lipid-rich microenvironment in CAP. Although FFPE or frozen section is standard procedure for histopathologic samples, lipid components are hardly preserved due to organic solvent exposure during those routine procedures. To validate LD in clinical samples of CAP, we obtained air-dried frozen sections and subjected them to Oil Red O staining. Surprisingly, we found exuberant LD in inflammatory cells as well as in extracellular space of CAP lesions, while those conventional H&E sections had a typical CAP histopathologically (Fig. 6e and Extended Data Fig. 10h). Taken together, our observations suggest an adaptive host-microbiome strategy in CAP whereby lipophilic CA persistently alters lipid metabolism within TREM2-M□ (Fig. 6f).

## Discussion

Although CAP is not life-threatening, more than 50% of the global adult population has at least one tooth affected^1^. CAP commonly develops following root canal (endodontic) treatment, with prevalence often ranging up to 20% in developed countries^39^. Although many populations of bacterial species are found in CAP^6,40,41^, apical surgery is mainly recommended due to the ineffectiveness of systemic or local antibiotics. In this study, we found the existence of CA in TREM2-M□ and a lipid-rich microenvironment in most CAP cases, explaining the therapeutic resistance of CAP. Considering that CA is commensal in skin, oral mucosa, and the GI tract, hindering its initial entry into the intrabony space (such as through trauma, dental caries or endodontic treatment) may be crucial in preventing CAP development.

In acne vulgaris, the most common skin disease worldwide, the pathologic role of CA is well-documented while effective therapeutics remain limited^42^. Although sebaceous gland or epithelium is largely regarded as a lipid-source for CA^20,30,31^, TREM2-mediated abundant LDs formation by CA suggest the possibility that innate immune cells are also lipid-providers during the acne vulgaris progression, especially during CA colonization and subsequent inflammatory stages. Furthermore, CA remains highly susceptible to routine antibiotics such as tetracyclines and beta-lactams, although topic application of those antibiotics are usually ineffective^43^. The lipid-rich environment induced by TREM2-M□ may provide a survival niche for CA, allowing it to escape from systemic or topical antibiotics. It should be noted that other immune cells, such as neutrophils and mast cells, can be lipid suppliers in inflammatory lesions, while we only focused on M□ in this study.

Gram positive anaerobic CA is predominantly found in acidic niches within pilosebaceous units of the skin, involving most common skin inflammation, acnes vulgaris. Interestingly, genetic analysis revealed that the genome and bacteriophage of CA have many homologs among mycobacteria and mycobacteriophages, respectively^44,45^. Based on similar histopathological features of tuberculosis and sarcoidosis, characterized by aggregation of activated macrophages in terms of “granuloma”, a Japanese group successfully identified CA as a potential etiologic link of sarcoidosis about fifty years ago^46^. Development of diagnostic antibodies for IHC and electron microscopic study revealed intracellular CA persistence in activated macrophages in clinical samples of sarcoidosis^47^. Interestingly, *Mtb* and CA are highly lipophilic bacterial species sharing the same host cells and a similar means of phagolysomal escape in M□^48^. While virulent *Mtb* can replicate within human M□^49^, we could not find evidence of CA replication in PBMC-M□, suggesting that low-virulent CA adapts a different strategy from *Mtb*. Interestingly, TREM2 has been identified as a receptor for mycolic acids in mycobacterium, anionic ligands of bacteria, and for lipo-polysaccharides of Niesseria^48,50,51^, suggesting that (1) CA also utilizes TREM2 for entry into M□, and (2) lipophilic ligands are present on the CA surface.

TREM2 is exclusively abundant in monocyte-macrophage cells, acting as a pivotal immune sensor for damaged cells and lipids^52^. As a cell surface lipid sensor, TREM2 regulates diverse functions, including lipid metabolism, by undergoing constitutive endocytosis and rapid trafficking to the cell surface^22,53^. In 2013, several groups reported TREM2 variants in Alzheimer’s, Parkinson’s and other neurodegerative diseases^16,17,54^. Subsequent large scale studies revealed the importance of TREM2 in various metabolic, degenerative, inflammatory diseases and tumor-associated M□ (TAM) in human cancer^55^. Loss of TREM2 in M□ impairs phagocytic activity and induces foamy macrophage by increased uptake of oxidized LDL^56^. Our study further reveals the importance of TREM2 in CA-mediated chronic inflammatory diseases in CAP and acne vulgaris^20^. Considering the prevalence of those diseases in the world, a deeper understanding of pathophysiology in CA-mediated chronic inflammation may significantly benefit human health.

In this study, we aimed to understand pathologic roles of intracellular persistent CA in CAP and the important role of TREM2-M□ in forming a lipid-rich microenvironment. In turn, CA has evolved an intriguing strategy involving TREM2-mediated LD formation to propagate as a human commensal. Further studies of CA-TREM2 interactions may elucidate uncharacterized pathophysiology of CAP in oral health as well as its implications for prosthetic inflammation, sarcoidosis and acne vulgaris.

## Methods

### Cell culture, transfection, and immunoblot analysis

Human peripheral blood mononuclear cells (PBMCs) were purchased from Goma Biotech (Lot No. 2207190101). Monocytes were isolated based on adherence by seeding PBMCs onto 100-mm culture dishes in monocyte attachment medium (PromoCell, C-28051) and incubating for 1 hour at 37°C in a humidified atmosphere with 5% CO□. After incubation, non-adherent cells were removed by gentle washing with phosphate-buffered saline (PBS), and the remaining adherent monocytes were cultured for differentiation. Monocyte-to-macrophage differentiation was induced by culturing the adherent cells in Dulbecco’s Modified Eagle Medium (DMEM) supplemented with 10% fetal bovine serum (FBS) and recombinant human granulocyte-macrophage colony-stimulating factor (GM-CSF; Gibco, #300-03) at a final concentration of 20 ng/mL for 6 days. The culture medium was refreshed every 3 days. Differentiated M□ were used for subsequent experiments.

THP-1 and RAW264.7 cells were obtained from Korean Cell Line Bank. AD293T cells were obtained from American Type Culture Collection (ATCC; Manassas, VA, USA). THP-1 cells were cultured in RPMI-1640, while RAW264.7 and AD293T cells were maintained in Dulbecco’s Modified Eagle Medium (DMEM), each supplemented with 10% fetal bovine serum (FBS) under standard culture conditions (37°C, 5% CO□).

### Generation of NLRP3 or TREM knock-out THP-1 cells

To generate NLRP3 or TREM2 knockout cells, lentiviruses from lentiCRISPRv2 (Addgene plasmid #52961 vectors)^57^ harboring sgRNAs targeting NLRP3 or TREM2 were transduced into THP-1 cells using spinfection methods (930g for 120 minutes at 30°C), followed by selection with puromycin (2 ug/ml). Single clones using limiting dilution were obtained for NLRP3 knockout, and pooled clones of lentiviral infected cells were used for TREM2. The loss of NLRP3 and TREM2 were examined by immunoblot analysis. CRISPR guide RNA sequences for lentiCRISPRv2 targeting NLRP3 and TREM2 were:

NLRP3 guide 1, 5’-GAGGGGTCAGACAGAGAAGG,

NLRP3 guide 2, 5’-GCTCACACTCTCACCCAGAC,

TREM2 guide 1, 5’-AGGGTGGCATGGAGCCTCTC,

TREM2 guide 2, 5’-TCTTACTCTTTGTCACAGgt (lowercase indicate intronic sequences).

### Generation of polyclonal antibody

10-week-old rabbits were acclimatized prior to immunization (Biocomplete, Korea). For initial immunization, live CA D1 isolate (2×10^9^ CFU) was injected subcutaneously in the flank. Four weeks after the initial injection, booster doses containing 1×10^9^ CFU of CA D1 were administered at 2-week intervals, for a total of four booster injections. Whole blood was collected in the SST (serum separating tube) and centrifuged at 3,000 rpm for 10 minutes to separate serum. The polyclonal antibody was purified from serum using a Protein A column affinity purification system. This antibody against D1 isolate was designated anti-CA antibody.

### CA culturing and preparation

CA were maintained and sub-cultured in thioglycollate broth. For experimental use, bacteria were sub-cultured into fresh brain-heart infusion (BHI) broth and incubated for 3-4 days to reach the logarithmic (proliferative) phase. All bacterial cultures were grown at 37°C under anaerobic conditions using the BD GasPak^TM^ system (Becton Dickinson). Bacterial density was estimated by measuring optical density at 600 nm (OD_600_) and converted to colony-forming units (CFU) using a previously established standard curve. Heat-killed *C. acnes* (D1) were prepared by incubation at 60°C for 30 minutes. Complete loss of viability was confirmed by the absence of colony formation on blood agar plate following inoculation. *Cutibacterium granulosum* (KCTC 5747) and *Cutibacterium avidum* (KCTC 5339) were obtained from the Korean Collection for Type Cultures (KCTC). These strains were cultured and handled under the same conditions as described for CA.

### Isolation and cultivation of CA clinical isolates

All procedures involving human samples were conducted in accordance with protocols approved by the Institutional Review Board of Yonsei University Dental Hospital (Yonsei University College of Dentistry, IRB No. 2-2022-0036), and written informed consent was obtained from all patients. The CA isolates CA-D1 and CA-D2 were isolated from peri-implantitis and apical periodontitis lesions, respectively. Tissue samples were enzymatically digested in Dulbecco’s Modified Eagle Medium (DMEM) containing type I collagenase (10 mg/mL; Gibco, #17100017) and DNase I (1 mg/mL; Roche, #10104159001) for 30 minutes at 37°C. The digested tissues were then passed through a cell strainer to obtain a single-cell suspension. Monocyte-lineage cells were enriched using CD14 MicroBeads (Miltenyi Biotec, #130-097-052) according to the manufacturer’s instructions. The isolated cells were cultured in DMEM supplemented with 2% penicillin/streptomycin/fungizone (P/S/F; Gibco, #15240062) for 24 hours, followed by cell lysis using a hypotonic solution. The resulting lysates were inoculated onto blood agar plates and incubated under anaerobic conditions to allow bacterial growth. Individual colonies were collected, and bacterial identity was confirmed as *C. acnes* by 16S rRNA gene PCR and sequencing using the following primers: forward 5′-GGGTTGTAAACCGCTTTCGCCT-3′ and reverse 5′-TTCGACGGCTCCCCCACAAC-3′.

### Single-locus sequence typing (SLST) analysis

Single-locus sequence typing (SLST) of *C. acnes* isolates was performed according to the method described by Christian F. Scholz et al.^26^. Briefly, the SLST target region was amplified by PCR using the following primers: forward 5′-CAGCGGCGCTGCTAAGAACTT-3′ and reverse 5′-CCGGCTGGCAAATGAGGCAT-3′. The PCR products were purified and subjected to Sanger sequencing using the same primer set. The obtained sequences were trimmed to the defined SLST region according to the reference scheme^26^ and aligned using the online SLST database (http://medbac.dk/slst/pacnes) to determine the corresponding SLST type.

### Antibiotic susceptibility testing

Antibiotic susceptibility of CA isolates was determined using a commercially available broth microdilution system (Sensititre™, Thermo Fisher Scientific) with anaerobic bacteria plates (ANO2B), according to the manufacturer’s instructions. Briefly, 3–5 colonies grown on blood agar plates were inoculated into brain–heart infusion (BHI) broth and cultured under anaerobic conditions for 72 hours. Bacterial density was then estimated by measuring optical density at 600 nm (OD_600_) and converted to colony-forming units (CFU). The bacterial suspension was diluted in Brucella broth to achieve a final inoculum of approximately 1 × 10□ CFU per 100 μL, and 100 μL of the suspension was dispensed into each well of the Sensititre plate. The plates were incubated under anaerobic conditions for 72 hours, and bacterial growth was assessed visually. The minimum inhibitory concentration (MIC) was defined as the lowest concentration of antibiotic at which no visible growth was observed. If bacterial growth was not observed at any tested concentration, the result was recorded less than the minimal concentration of the plate.

### Immunofluorescence (IF) study

Cells were fixed with 4% paraformaldehyde (PFA) for 10 minutes at room temperature and permeabilized with 0.5% Triton X-100 in phosphate-buffered saline (PBS) for 10 minutes. After washing, cells were blocked with 5% normal goat serum for 1 hour at room temperature. Primary antibodies were applied and incubated overnight at 4°C. Following incubation, cells were washed with PBS containing 0.1% Tween-20 (PBS-T) and incubated with appropriate Alexa Fluor-conjugated secondary antibodies (Invitrogen; Alexa Fluor 488, 594, and 647) at a dilution of 1:1,000 for 1 hour at room temperature in the dark. After washing, nuclei were counterstained and mounted using ProLong™ Gold Antifade Mountant with DAPI (Invitrogen). Fluorescence images were acquired using Zeiss LSM700 or LSM780 confocal laser scanning microscopes.

### Transmission Electron Microscopy (TEM)

RAW 264.7 cells and hPBMC-derived macrophages were infected with CA-D1 or HK-CA at a multiplicity of infection (MOI) of 100 and 50, respectively. Cells were fixed at 1, 3, and 5 days post-infection period for 24 hours in a solution containing 2% glutaraldehyde and 2% paraformaldehyde in 0.1 M phosphate buffer (pH 7.4), followed by post-fixation in 1% osmium tetroxide (OsO□) for 2 hours. Samples were dehydrated through a graded ethanol series (50%–100%) and infiltrated with propylene oxide for 10 minutes. The specimens were then embedded in Poly/Bed 812 resin (Polysciences) and polymerized at 70°C for 12 hours. Semithin sections (200 nm) were cut using an ultramicrotome (UC7), stained with toluidine blue, and examined under a light microscope. Ultrathin sections (80 nm) were collected on copper grids, stained with 5% uranyl acetate for 20 minutes and 3% lead citrate for 7 minutes. Images were acquired using a Hitachi HT7800 transmission electron microscope equipped with a digital camera, operated at an accelerating voltage of 100 kV.

### Cytokine array analysis

Cytokine profiling was performed using the RayBio® C-Series Human Inflammation Antibody Array C3 (RayBiotech, #AAH-INF-3-8) according to the manufacturer’s instructions. Briefly, PBMC-M□ were infected with CA at a multiplicity of infection (MOI) of 20. After 24 hours of incubation, culture supernatants were collected and applied to the antibody array membranes. Signal detection was performed following the manufacturer’s protocol, and chemiluminescent signals were visualized using X-ray film. Relative cytokine expression levels were analyzed by densitometric quantification.

### IHC and Multiplex IHC

Seven FFPE CAP tissues were collected for pathological and IHC examination. The H&E sections were examined by pathology specialists (HSK and JIY). IHC analysis was performed to investigate the presence of intracellular infection of CA in CAP tissue. Five μm thickness tissue sections were deparaffinized and rehydrated followed by antigen retrieval in 10 mM sodium citrate buffer (pH 6.0) at 94^0^C for 30 min. Endogenous peroxidase activity was blocked with 3% H2O2 solution, followed by blocking and permeabilization in PBS containing 2.5% normal goat serum and 0.3% Triton X-100. Sections were then incubated overnight at 4^0^C with primary antibody (PAB antibody 1:10 000, MBL Life Science) specific for CA. After washing, sections were incubated for 1 h with HRP-conjugated anti-mouse/rabbit IgG and developed with diaminobenzidine (DAB) (Agilent, K500711-2). Counterstaining was performed with Mayer’s hematoxylin.

For mIHC, FFPE samples were stained for CD68 (Agilent, Cat. No. M0876), CD14 (Abcam, Cat. No. ab183322) and PAB (MBL Life Science, Cat. No. D371-3) using the TSA-Opal method on the Leica BOND RX automated immunostainer (Leica Microsystems). All tissue samples were deparaffinized and subjected to antigen retrieval using BOND Epitope Retrieval 1 (ER1) (Leica Biosystems, Cat. No. AR9961), then treated following this protocol: all incubation steps were performed at room temperature (RT), then followed by four 1-minute washes using BOND Wash Solution (Leica Biosystems, Cat. No. AR9590). The automated protocol began with a protein block using 1X Antibody Diluent/Block (Quanterix, Cat. No. ARD1001EA) for 10 min at RT. Tissues were incubated with the anti-CD68 antibody (Agilent, Cat. No. M0876) for 60 min. The slides were then incubated with the 1X Opal Anti-Ms+Rb HRP (detection polymer, Quanterix, Cat. No. ARH1001EA) for 20 min. Opal 570 Reagent (Quanterix, Cat. No. FP1488001KT) was applied at a 1:150 dilution for 10 min at RT. For the second cycle, the bound antibody stripping was performed using BOND Epitope Retrieval 2 (ER2) (Leica Biosystems, Cat. No. AR9640) by heating the slides at 95□ for 20 min. Following stripping, the protocol was repeated for anti-CD14 primary antibody (Abcam, Cat. No. ab183322) detecting Opal 520 reagent and anti-PAB primary antibody (MBL Life Science, Cat. No. D371-3) detecting Opal 690 reagent. After the final Opal staining, nuclear counterstaining was performed using DAPI (Quanterix, Cat. No. FP1490) for 10 min at RT. All stained slides were digitized using the PhenoImager HT (formerly Vectra Polaris) mutispectral imaging system (Quanterix). Multispectral images were acquired at 40x magnification to capture the fluorescent signal from all applied fluorophores. The specific channels acquired included DAPI (Nuclear Counterstain), Opal 570 (CD68 marker), Opal 520 (CD14 marker) and Opal 690 (PAB marker). Following image acquisition, regions of interest (ROIs) suitable for analysis were designated using PhenoChart software (Quanterix). Cells within the ROIs were then analyzed using inForm software (Quanterix) through the following sequential steps: spectral unmixing to remove autofluorescence, followed by cell segmentation and phenotyping based on staining patterns. The established algorithm was applied uniformly across all tissue samples.

### Bulk RNA sequencing and analysis

Bulk RNA sequencing was performed following total RNA extraction with Trizol reagent (Invitrogen. #15596018). Paired-end sequencing (100 bp) was conducted on the NextSeq 550 platform (Novogene, China). Raw sequencing reads were processed using fastp to remove adapter sequences and low-quality reads. The resulting high-quality reads were aligned to the human reference genome (GRCh38, ensembl_100) using HISAT2 (v2.2.1). Gene-level read counts were subsequently quantified using featureCounts (v2.0.6). All statistical analyses were performed in the R statistical environment (v4.4.2). For differential expression analysis between the infected and control groups, only protein-coding genes were retained, and low-expression genes were filtered out. The raw read counts were normalized using the Trimmed Mean of M-values (TMM) method. Differentially expressed genes (DEGs) were identified using a generalized linear model (GLM) likelihood ratio test implemented in the edgeR package (v4.4.2). Statistical significance was defined by an absolute log2 fold change > 1 and a False Discovery Rate (FDR) < 0.05. For visualization, normalized counts were converted to log2-transformed Transcripts Per Million (TPM). Unsupervised hierarchical clustering of the DEGs was performed using Pearson correlation as the distance metric and Ward’s linkage method. Additionally, volcano plots were generated to visualize the magnitude of gene expression changes alongside their statistical significance. Furthermore, to assess sample relationships, pairwise scatter plots displaying Pearson correlation coefficients were generated. A sample correlation matrix was also visualized using the pheatmap package (v1.0.13) based on the log2-transformed TPM values. To elucidate the biological functions and pathways associated with the DEGs, over-representation analysis (ORA) for Gene Ontology (GO) terms and Kyoto Encyclopedia of Genes and Genomes (KEGG) pathways was performed using the clusterProfiler package (v4.14.6). ENSEMBL gene IDs were mapped to Entrez IDs via the org.Hs.eg.db database (v3.20.0), and all expressed genes within the dataset were designated as the background universe. Statistical significance of the enrichment was evaluated using the Benjamini-Hochberg procedure, with an FDR < 0.05 threshold. To comprehensively visualize the top 25 significantly enriched pathways, a dual-axis plot was generated to concurrently display the statistical significance of the enrichment (-log10 FDR) and the exact percentage of up-regulated genes within each enriched term, providing deeper functional insights into the transcriptional alterations.

### *In vivo* experiments

Seven-week-old male C57bl/6 mice were used for subcutaneous injection experiments following a one-week acclimation period. The dorsal skin was shaved using a trimmer and disinfected with 70% ethanol swabs prior to injection. Live *C. acnes* (D1 or D2 isolates) and heat-killed *C. acnes* were suspended in a hyaluronic acid and PBS mixture (6:4 ratio) to enhance local retention. A total of 1×10^8^ CFU in 100μL volume was injected subcutaneously into both sides of the dorsal regions. Mice received two injections at the same sites at 7 days apart and were euthanized 7 days after the final injection. Skin tissues from the injection sites, including dermis to the depth of skeletal muscle tissue or peritoneum, were harvested and immediately fixed in 10% neutral buffered formalin for 24 hours. Fixed tissues were processed for paraffin embedding (FFPE) and sectioned for histological and immunohistochemical analysis. The animal study scheme was approved by the Institutional Animal Care and Use Committee (IACUC) of Yonsei University College of Medicine (IACUC No.2024-0145).

### Enzyme-linked immunosorbent assay (ELISA)

Human IL-1β levels in culture supernatants were measured using a commercially available ELISA kit (Invitrogen, KHC0011) according to the manufacturer’s instructions. THP-1 cells were infected with *C. acnes* (CA-D1) at MOI 100, while PBMC-M□ were infected at MOI 20. After 24 h, culture supernatants were collected and directly subjected to ELISA without dilution. Absorbance was measured at 450 nm using a microplate reader, and cytokine concentrations were calculated based on a standard curve generated according to the manufacturer’s protocol.

### Human CAP samples and scRNA-seq analysis

The procedures for the collection of CAP samples at the Department of Conservative Dentistry of Yonsei University Dental Hospital was approved by the Ethic Commission of the Dental Hospital (IRB No. 2-2024-0097) and the patients gave their written informed consent. Periapical surgery of selected CAP cases was performed by a professional endodontist (D.H.K.). The CAP samples were collected immediately in ice-cold DMEM prior to processing. Isolated single cells were loaded onto a microwell cartridge of the BD Rhapsody Express HT system (BD) according to the manufacturer’s instructions. Single cell capture and mRNA capture were performed using the BD Rhapsody Cartridge kit (633733; BD), Cartridge Enhanced Reagent kit (664887; BD), and BD Rhapsody cDNA kit (633773; BD). Whole-transcriptome libraries were subsequently generated using the BD Rhapsody WTA Reagent Kit (633802; BD) following the manufacturer’s instructions. The final indexed libraries were sequenced on an Illumina NextSeq2000 platform using the P3 reagent kit (100 cycles; Illumina). Sequencing was performed with 50 + 75 bp paired-end reads and an 8-bp single index, targeting a sequencing depth of more than 20,000 reads per cell for each sample. The FASTQ-format sequencing raw data was initially processed using the BD Rhapsody Sequencing Analysis Pipeline (version 1.2; VELSERA). Reads were aligned to the hg38 human reference genome using the BD Rhapsody reference package (RhapRef_Human_WTA_2025-03). Data normalization, dimensionality reduction and visualization were performed in R using the Seurat package (version 4.4.0), unless otherwise specified. Raw gene expression matrices were filtered to remove low-quality cells. Cells with a high proportion of mitochondrial gene expression (>60%), detection of ambient hemoglobin gene expression (HBA1, HBA2 and HBA2), or evidence of doublets/multiplets were excluded from downstream analyses. Filtered datasets from CAP lesions and healthy control tissues were normalized and log-transformed using LogNormalize with a scale factor of 10,000. Highly variable genes were identified using the vst method (2,000 genes/sample) and used for downstream data integration and dimensionality reduction. Samples were integrated using FindIntegrationAnchors and IntegrateData with dimensions 1–20. The integrated assay was scaled with regression of total UMI counts and mitochondrial transcript percentage, followed by principal component analysis, shared nearest neighbor graph construction, clustering, and UMAP visualization. Clustering resolution was selected empirically after evaluation across multiple resolutions, and marker genes were identified using FindAllMarkers (only.pos = TRUE, min.pct = 0.25, logfc.threshold = 0.25). Cell populations were annotated manually based on canonical marker gene expression. For myeloid subclustering analysis, myeloid cells were subsetted and re-analyzed using the same workflow, including normalization, scaling, PCA, clustering, and UMAP visualization. Gene set expression scores were calculated using module scoring implemented in Seurat (AddModuleScore). To quantify lipid metabolic programs, curated gene sets representing distinct lipid metabolism pathways were manually compiled based on prior literature and pathway annotations. These included fatty acid transport and uptake, fatty acid oxidation (FAO), lipogenesis/fatty acid synthesis, triglyceride synthesis, lipid droplet storage, lipolysis, cholesterol biosynthesis, cholesterol efflux and trafficking, phospholipid remodeling, sphingolipid/ceramide metabolism, arachidonic acid/eicosanoid metabolism, and core lipid regulatory programs. Gene lists for each lipid metabolism pathway are provided in Supplementary Table 4. Pseudotime analysis was conducted using Slingshot (version 2.10.0), and cells were ordered along the inferred trajectory. Gene expression dynamics along pseudotime were visualized using heatmaps and smoothed trend curves.

### Statistical analysis

Statistical analyses were performed using GraphPad Prism 8 (GraphPad Software, San Diego, CA, USA). Comparisons between two groups were conducted using the Mann–Whitney U test, while comparisons among multiple groups were performed using the Kruskal–Wallis test followed by Dunn’s multiple comparisons test for post hoc analysis against the control group. Statistical significance was defined as p < 0.05 (*), < 0.01 (**), and < 0.001 (***).

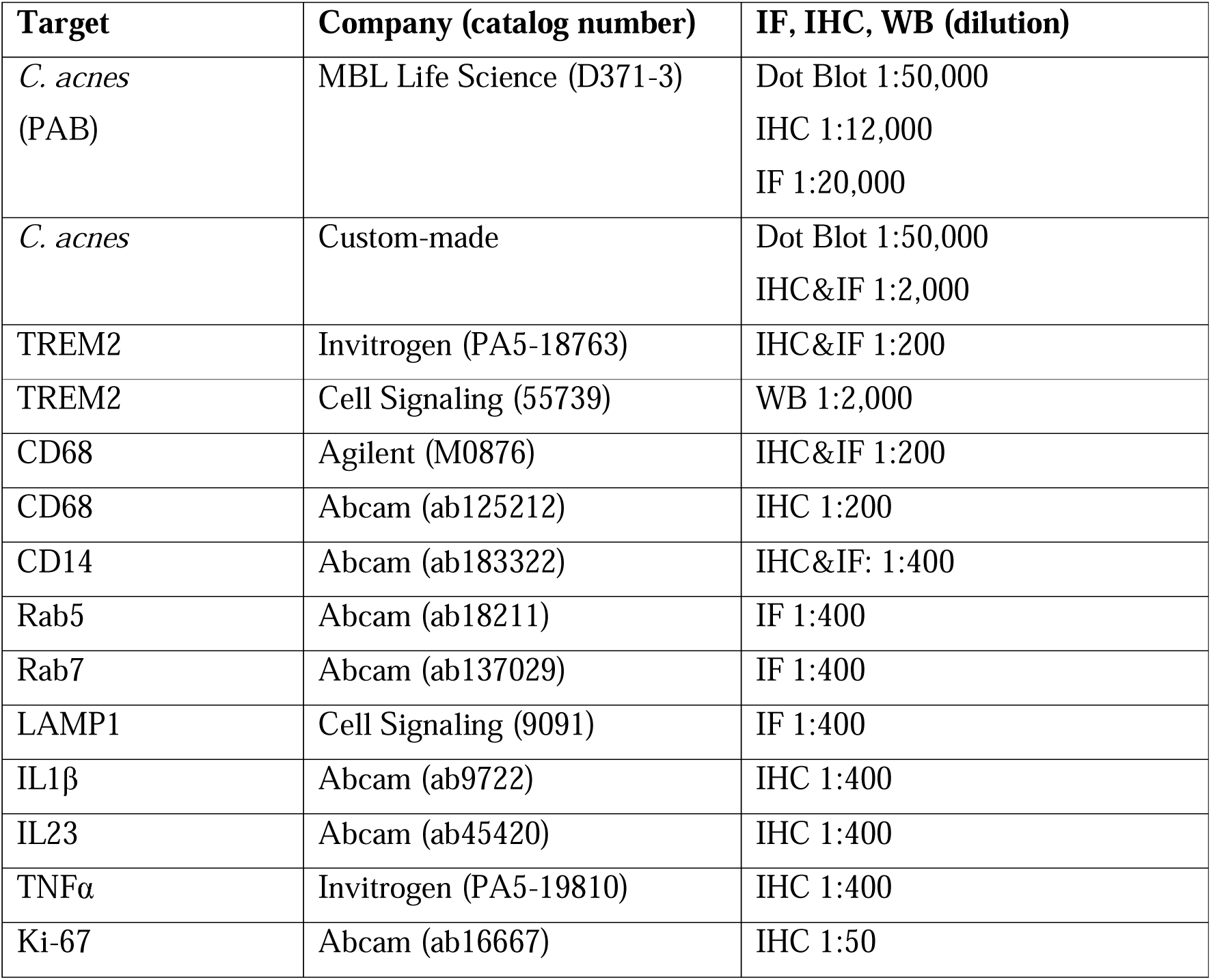
Antibodies used in this study.

Antibodies used in this study
| Target | Company (catalog number) | IF, IHC, WB (dilution) |
| --- | --- | --- |
| *C. acnes*<br>(PAB) | MBL Life Science (D371-3) | Dot Blot 1:50,000<br>IHC 1:12,000<br>IF 1:20,000 |
| *C. acnes* | Custom-made | Dot Blot 1:50,000<br>IHC&IF 1:2,000 |
| TREM2 | Invitrogen (PA5-18763) | IHC&IF 1:200 |
| TREM2 | Cell Signaling (55739) | WB 1:2,000 |
| CD68 | Agilent (M0876) | IHC&IF 1:200 |
| CD68 | Abcam (ab125212) | IHC 1:200 |
| CD14 | Abcam (ab183322) | IHC&IF: 1:400 |
| Rab5 | Abcam (ab18211) | IF 1:400 |
| Rab7 | Abcam (ab137029) | IF 1:400 |
| LAMP1 | Cell Signaling (9091) | IF 1:400 |
| IL1β | Abcam (ab9722) | IHC 1:400 |
| IL23 | Abcam (ab45420) | IHC 1:400 |
| TNFα | Invitrogen (PA5-19810) | IHC 1:400 |
| Ki-67 | Abcam (ab16667) | IHC 1:50 |

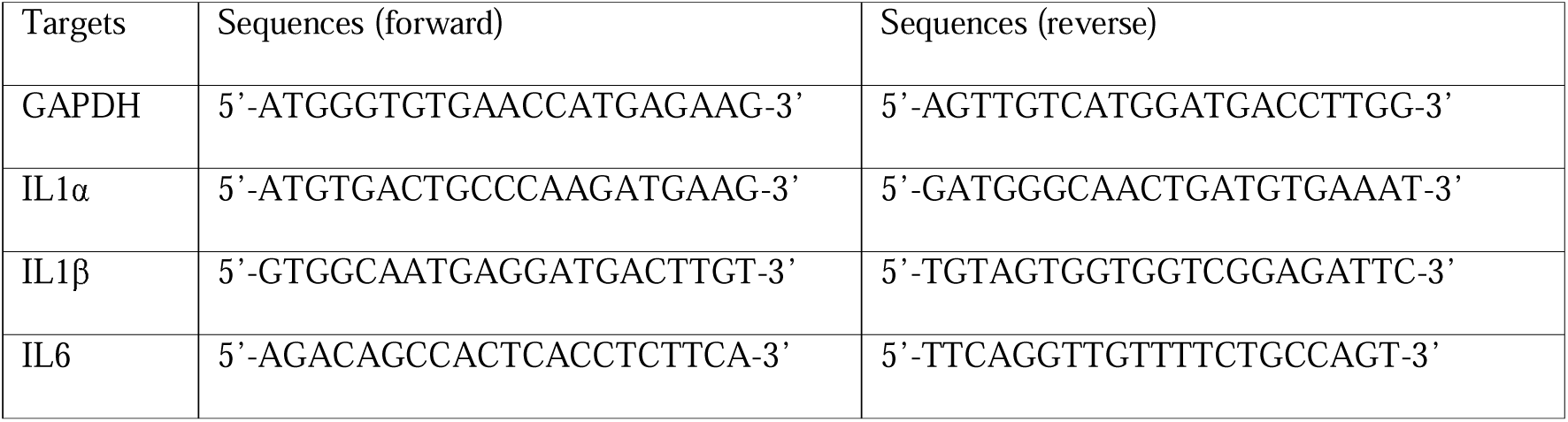
PCR primers.

## AUTHOR CONTRIBUTIONS

D.H. and N.H.K. performed main experiments; S.Y.H., H.E.S., S.Y.C., Y.C.B., K.Y.K., S.R.K., supported in vitro and in vivo experiments; S.B.C. performed bioinformatics analysis; H.K., J.Y.P., and J.S.L. provide clinical samples of peri-implantitis; S.J.S. advised research plan, S.L., performed mIHC analysis; E.S.C., H.S.K., D.H.K., and J.I.Y. planned experiments, analyzed the data, and wrote the manuscript.

## ACKNOWLEDGMENTS

We thank E. Tunkle for preparation of the manuscript and Jun Cheol Hong for technical assistance. We also thank Dr. Kwangwon Hong (Biocomplete, Korea) for generation of polyclonal antibody and Dr. Hyun-Woo Jeong (Einocle, Korea) for scRNA analysis of clinical samples.

## Supplementary Video 1-3

To obtain EGFP expressing *E. coli*, the BL21(DE3) was transformed with EGFP expressing pET-24a plasmid. A single colony was grown at 25°C until mid-log phase (OD_600_ 0.4-0.6). Then, 0.5 mM IPTG (isopropyl-D-1-thiogalactopyranoside) was added to induce EGFP overnight. The live *E. coli* (MOI 20) were added to the cell lines to obtain live images (1 min interval for hPBMC-M□ and THP-1, 4 min interval for RAW264.7) using Juli^TM^ stage (NanoEnteck).

