## supplemental figures for "Interaction between TREM2-Macrophages and *Cutibacterium acnes* Drives Altered Lipid Metabolism in Chronic Apical Periodontitis"

**Extended Data Fig. 1**

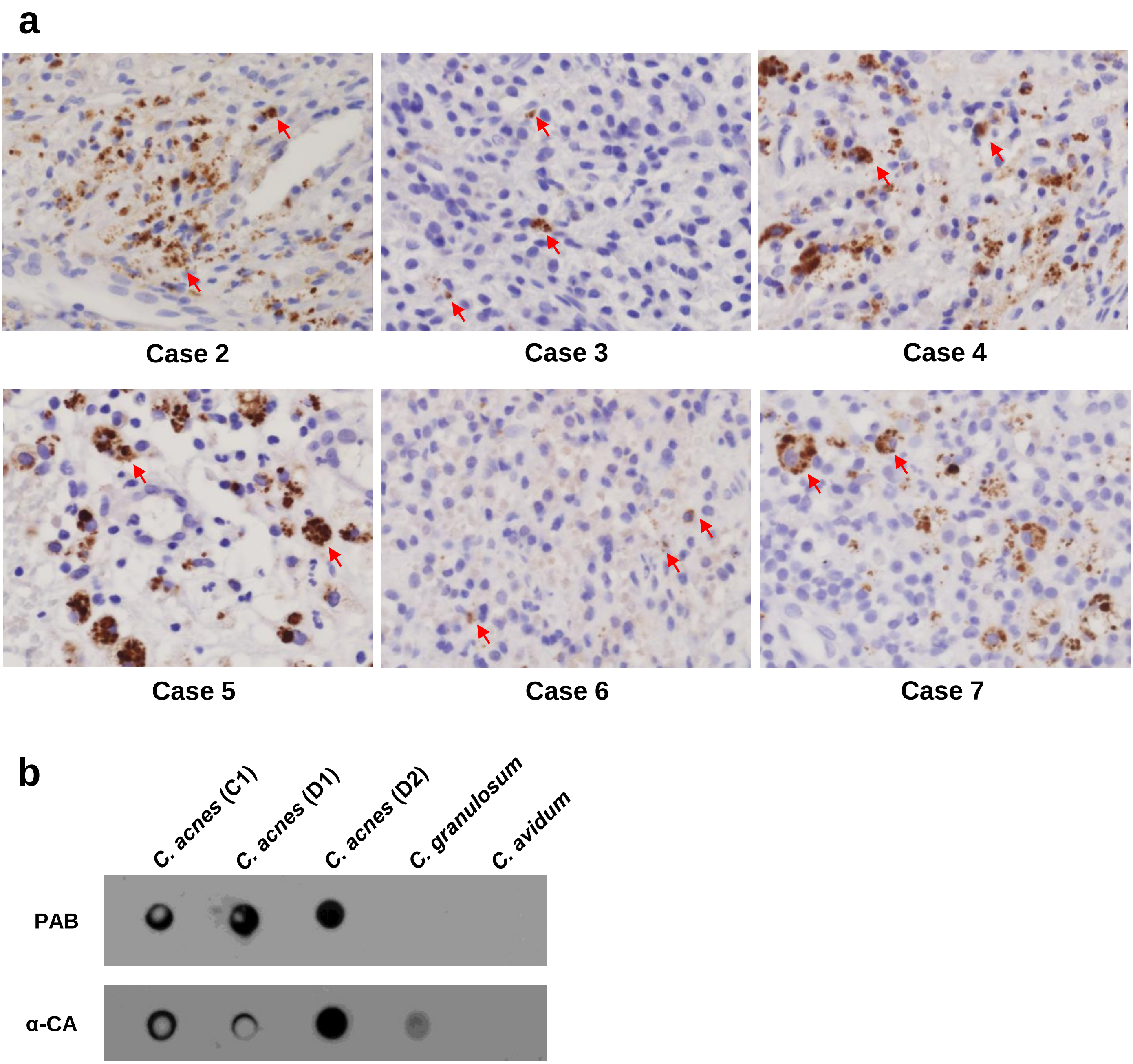


**Extended Data Fig. 1. CA persistence in macrophages in CAP lesions. a**. Detection of intracellular CA in CAP samples by IHC. Representative IHC staining of FFPE tissues from additional CAP cases (Cases 2-7) using a PAB antibody. PAB-positive CA signals were observed as punctate cytoplasmic staining within inflammatory cells (red arrows), consistent with intracellular localization across 7 cases. **b**. Reactivity and sensitivity of PAB and anti-*C. acnes* (α-CA) antibodies assessed by dot blot analysis. Dot blot analysis was performed using CA-D1, D2, C1 strains and related species (*C. granulosum* and *C. avidum*). Bacterial suspensions ($2 \times 10^7$ CFU in 20 μL) were spotted onto nitrocellulose membranes and probed with PAB or α-CA antibodies (1:50,000 dilution). Both PAB and α-CA antibodies showed strong reactivity toward all CA strains tested. While PAB exhibited minimal cross-reactivity with other *Cutibacterium* species, α-CA showed weak reactivity with *C. granulosum*.

### Extended Data Fig. 2.

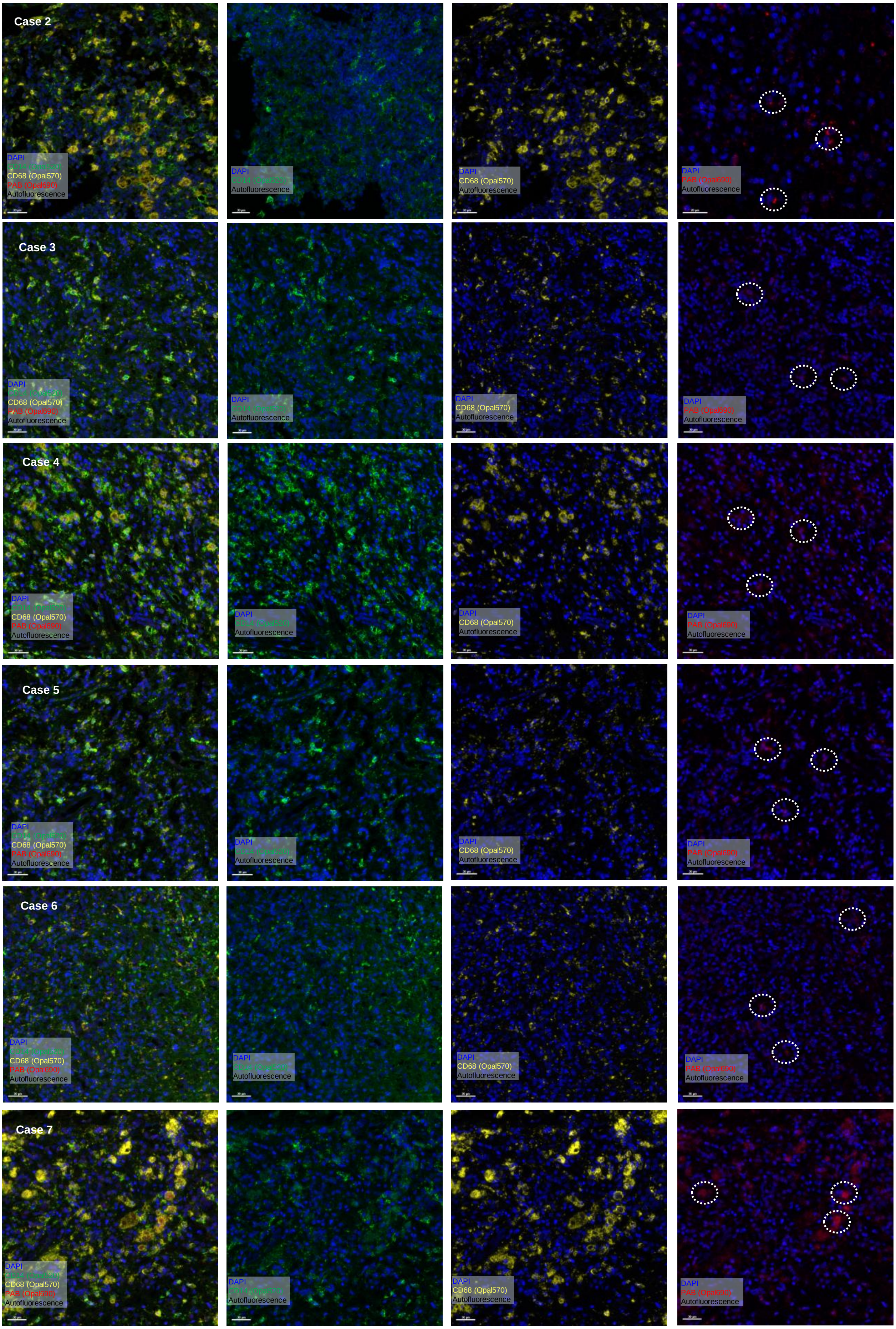

**Extended data Fig. 2. Representative images of mIHC analysis of CA localization in CAP tissues**. mIHC was performed on FFPE tissues from additional CAP cases using antibodies against CD14, CD68, and PAB. Representative images from multiple cases demonstrate CA signals co-localized with $CD14^{+}$ and/or $CD68^{+}$ cells (white dotted circles), consistent with intracellular localization within inflammatory cells.

**Extended Data Fig. 3.**

**a**

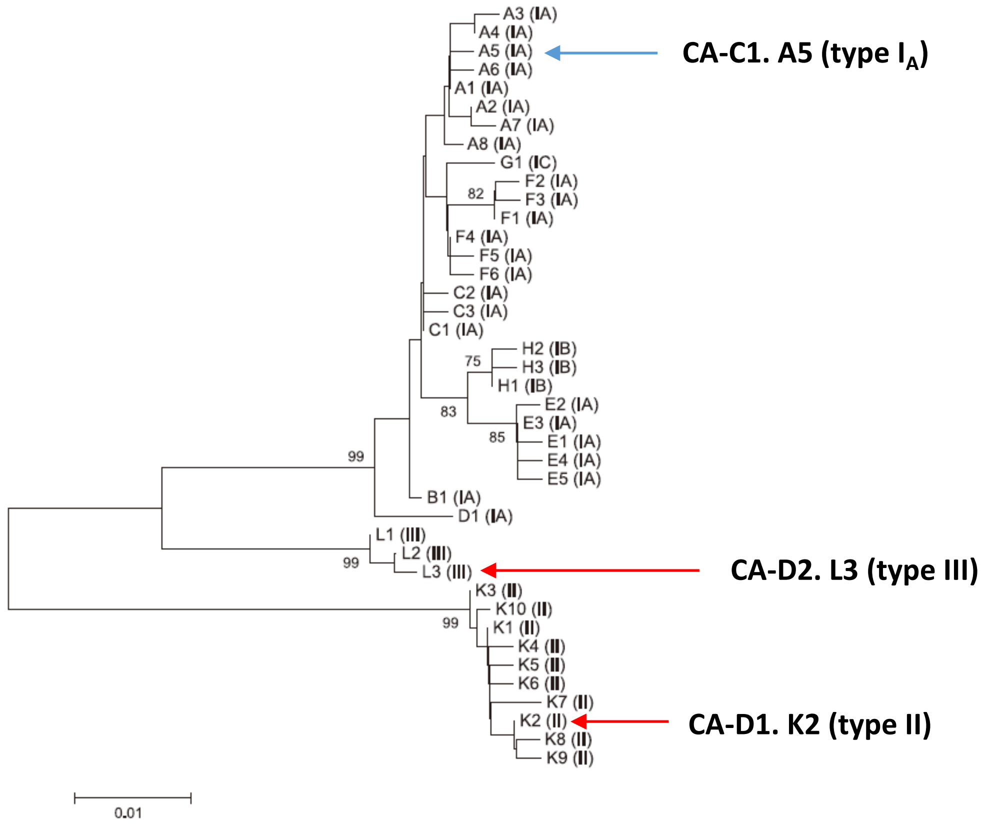


**b**

| Antibiotics | MIC50 | MIC90 | Range of plate | CA-D1 | CA-D2 |
|---|---|---|---|---|---|
| Ampicillin/sulbactam (2:1) | 0.25/0.06 | | 0.5/0.25 ~ 16/8 | <0.5/0.025 | <0.5/0.025 |
| Amoxicillin/clavulanic acid (2:1) | 0.25 | 0.50 | 0.5/0.25 ~ 16/8 | <0.5/0.025 | <0.5/0.025 |
| Cefotetan Na | n/a | n/a | 4 ~ 64 | <4 | <4 |
| Penicillin | 0.0625 | 0.125 | 0.06 ~ 4 | 0.12 | <0.06 |
| Imipenem | n/a | n/a | 0.12 ~ 8 | <0.12 | <0.12 |
| Meropenem | 0.06 | 1.5 | 0.5 ~ 8 | <0.5 | <0.5 |
| Clindamycin | 0.032 | 8.0 | 0.25 ~ 8 | <0.25 | <0.25 |
| Cefoxitin | 4 | | 1 ~ 32 | <1 | <1 |
| Metronidazole | >32 | | 0.5 ~16 | >16 | >16 |
| Chloramphenicol | 3.1 | | 2~64 | 8 | <2 |
| Ampicillin | 0.0312 | 0.125 | 0.5 ~ 16 | <0.5 | <0.5 |
| Piperacillin | n/a | n/a | 4 ~ 128 | <4 | <4 |
| Tetracycline | 0.19 | 4 | 0.25 ~8 | 2 | 1 |
| Mezlocillin | n/a | n/a | 4 ~ 128 | <4 | <4 |
| Piperacillin/tazobactam constant4 | 1/4 | | 0.25/4 ~ 128/4 | 2/4 | <0.25/4 |

**Antibiotics Susceptibility Test of CA-D1, D2 (Sensititre)** **(μg/mL)**

**Extended data Fig. 3**. **Phylogenetic analysis of CA isolates and antibiotic susceptibility**. **a**. Phylogenetic classification of clinical CA isolates based on SLST typing. Phylogenetic tree of CA phylotypes adapted from Christian F. Scholz et al.26. The positions of clinical isolates used in this study are indicated: CA-C1 (SLST type A5, type IA), CA-D1 (SLST type K2, type II), and CA-D2 (SLST type L3, type III). The tree illustrates the relationship between SLST types and classical phylogenetic classification of CA. Figure reproduced under CC BY license. **b**. Antibiotic susceptibility profiles of clinical CA isolates. Antibiotic susceptibility of CA-D1 and CA-D2 was determined using the Sensititre™ anaerobic bacteria panel (ANO2B). Minimum inhibitory concentrations (MICs) are shown for each antibiotic. $MIC_{50}$ and $MIC_{90}$ values were compiled from previously published studies[8,28,58-60]. All concentrations are expressed in μg/mL. For most antibiotics, no bacterial growth was observed at the lowest concentration tested, and values are therefore presented as below the lower detection limit (<). Consistent with previous reports, both isolates exhibited resistance to metronidazole and chloramphenicol.

**Extended Data Fig. 4.**

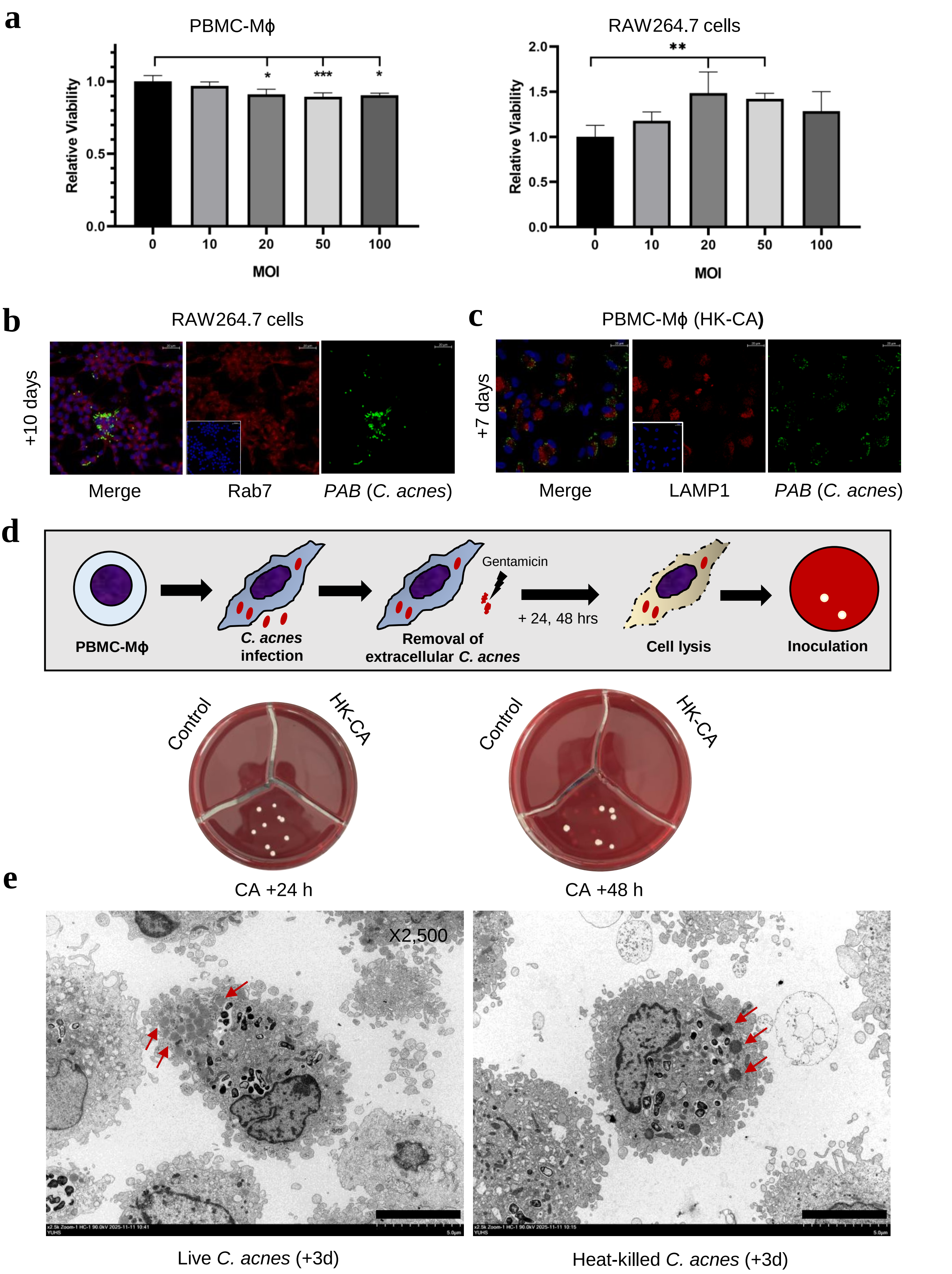

**Extended Data Fig. 4**. **Low-virulent intracellular persistence of CA in Mϕ**. **a**. Effects of CA infection on macrophage viability. hPBMC-Mϕ and RAW 264.7 cells were infected with CA at the indicated multiplicities of infection (MOIs). Cell viability was assessed using an MTT assay. Overall, CA infection resulted in only modest changes in cell viability across the tested MOI range in RAW264.7 cells, and in a slight reduction observed in hPBMC-Mϕ. **b**. Persistent intracellular localization of CA in RAW 264.7 cells. Representative IF images of RAW 264.7 cells infected with CA and analyzed at 10 days post-infection. CA signals (green) were detected within cells and showed limited co-localization with the late endosomal marker Rab7 (red). Intracellular bacterial signals remained detectable over time, consistent with prolonged intracellular persistence. (Scale bars, 20 μm) **c**. Intracellular persistence of heat-killed CA (HK-CA) in Mϕ. Representative IF images of hPBMC-Mϕ treated with HK-CA and analyzed at 7 days post-treatment. HK-CA signals (green) were detected within cells, with partial overlap with the lysosomal marker LAMP1 (red). Intracellular signals remained detectable over time, suggesting that prolonged intracellular retention may not be solely dependent on bacterial viability. (Scale bars, 20 μm) **d**. Recovery of viable intracellular CA from infected macrophages. Schematic overview (upper panel) and representative colony formation assay of hPBMC-Mϕ infected with CA (MOI 40). To eliminate extracellular bacteria, cells were treated with gentamicin (200 μg/mL) after infection. At 24 and 48 hours post-infection, macrophages were lysed and cytosolic fractions were plated under anaerobic conditions. Colonies were recovered from macrophages infected with live CA-D1, indicating that intracellular bacteria remained viable and retained the capacity to replicate on agar plates. In contrast, no colonies were detected in cells treated with HK-CA. **e**. Lipid droplets (LD) accumulation in Mϕ by persistent CA. TEM images of hPBMC-Mϕ exposed to live or HK-CA and analyzed at 3 days post-infection. Both live and HK-CA were associated with increased lipid droplet structures in the cytoplasm of Mϕ (red arrows).

### Extended Data Fig. 5. macrophage response (RAW RNA seq) and NLRP3-dependent inflammasome activation

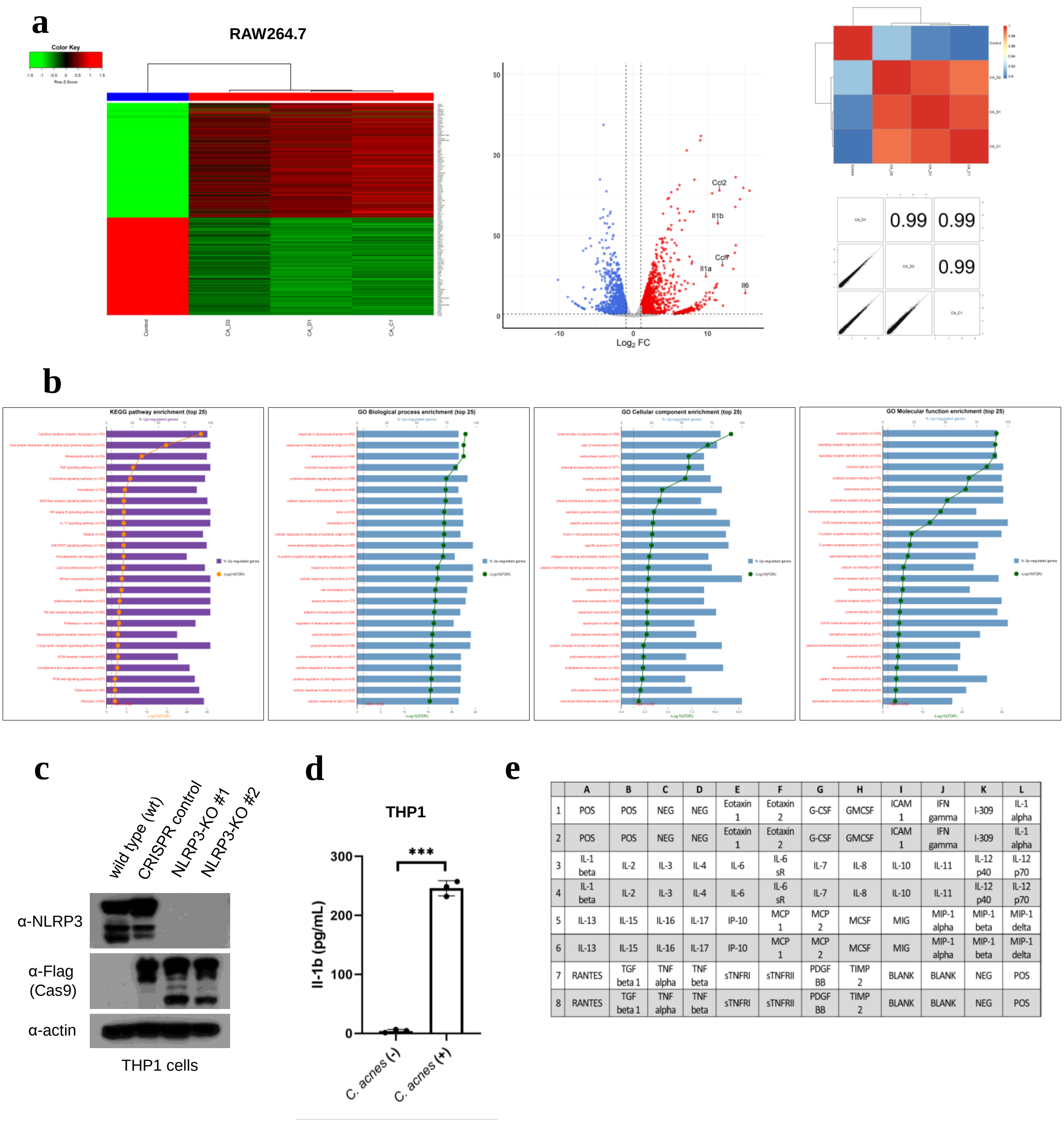


| | A | B | C | D | E | F | G | H | I | J | K | L |
|---|---|---|---|---|---|---|---|---|---|---|---|---|
| 1 | POS | POS | NEG | NEG | Eotaxin 1 | Eotaxin 2 | G-CSF | GMCSF | ICAM 1 | IFN gamma | I-309 | IL-1 alpha |
| 2 | POS | POS | NEG | NEG | Eotaxin 1 | Eotaxin 2 | G-CSF | GMCSF | ICAM 1 | IFN gamma | I-309 | IL-1 alpha |
| 3 | IL-1 beta | IL-2 | IL-3 | IL-4 | IL-6 | IL-6 sR | IL-7 | IL-8 | IL-10 | IL-11 | IL-12 p40 | IL-12 p70 |
| 4 | IL-1 beta | IL-2 | IL-3 | IL-4 | IL-6 | IL-6 sR | IL-7 | IL-8 | IL-10 | IL-11 | IL-12 p40 | IL-12 p70 |
| 5 | IL-13 | IL-15 | IL-16 | IL-17 | IP-10 | MCP 1 | MCP 2 | MCSF | MIG | MIP-1 alpha | MIP-1 beta | MIP-1 delta |
| 6 | IL-13 | IL-15 | IL-16 | IL-17 | IP-10 | MCP 1 | MCP 2 | MCSF | MIG | MIP-1 alpha | MIP-1 beta | MIP-1 delta |
| 7 | RANTES | TGF beta 1 | TNF alpha | TNF beta | sTNFRI | sTNFRII | PDGF BB | TIMP 2 | BLANK | BLANK | NEG | POS |
| 8 | RANTES | TGF beta 1 | TNF alpha | TNF beta | sTNFRI | sTNFRII | PDGF BB | TIMP 2 | BLANK | BLANK | NEG | POS |

**Extended Data Fig 5. Acute inflammatory transcriptional and cytokine response of CA infected macrophages**. **a**. Comparative transcriptomic responses to different CA isolates in Mϕ. RAW 264.7 cells were infected with CA isolates (CA-C1, CA-D1, and CA-D2), followed by bulk RNA sequencing analysis. Heatmap (left panel) and volcano plot (middle panel) analyses revealed similar transcriptional responses across all isolates, with consistent upregulation of pro-inflammatory genes, including Il1α, Il1β, Il6, and Ccl2. Correlation analysis further demonstrated high similarity in gene expression profiles among those isolates (right panels). **b**. Pathway enrichment analysis of CA–induced transcriptional responses. KEGG and Gene Ontology (GO) enrichment analyses were performed on differentially expressed genes identified in CA-infected hPBMC-Mϕ. Upregulated genes were enriched in inflammatory and immune-related pathways, including cytokine signaling and TNF signaling, consistent with the pro-inflammatory transcriptional profile observed in Fig. 3a. **c**. Western blot analysis confirming NLRP3 knockout in THP1 cells using CRISPR/Cas9-mediated gene editing. Expression of NLRP3, Cas9 (Flag), and β-actin is shown. **d**. ELISA analysis of IL-1β in CA infected THP1 cells. Wild-type THP1 cells were infected with CA (MOI 100) and IL-1β levels in culture supernatants measured.

**e**. Layout of the cytokine array panel.

**Extended Data Fig. 6.**

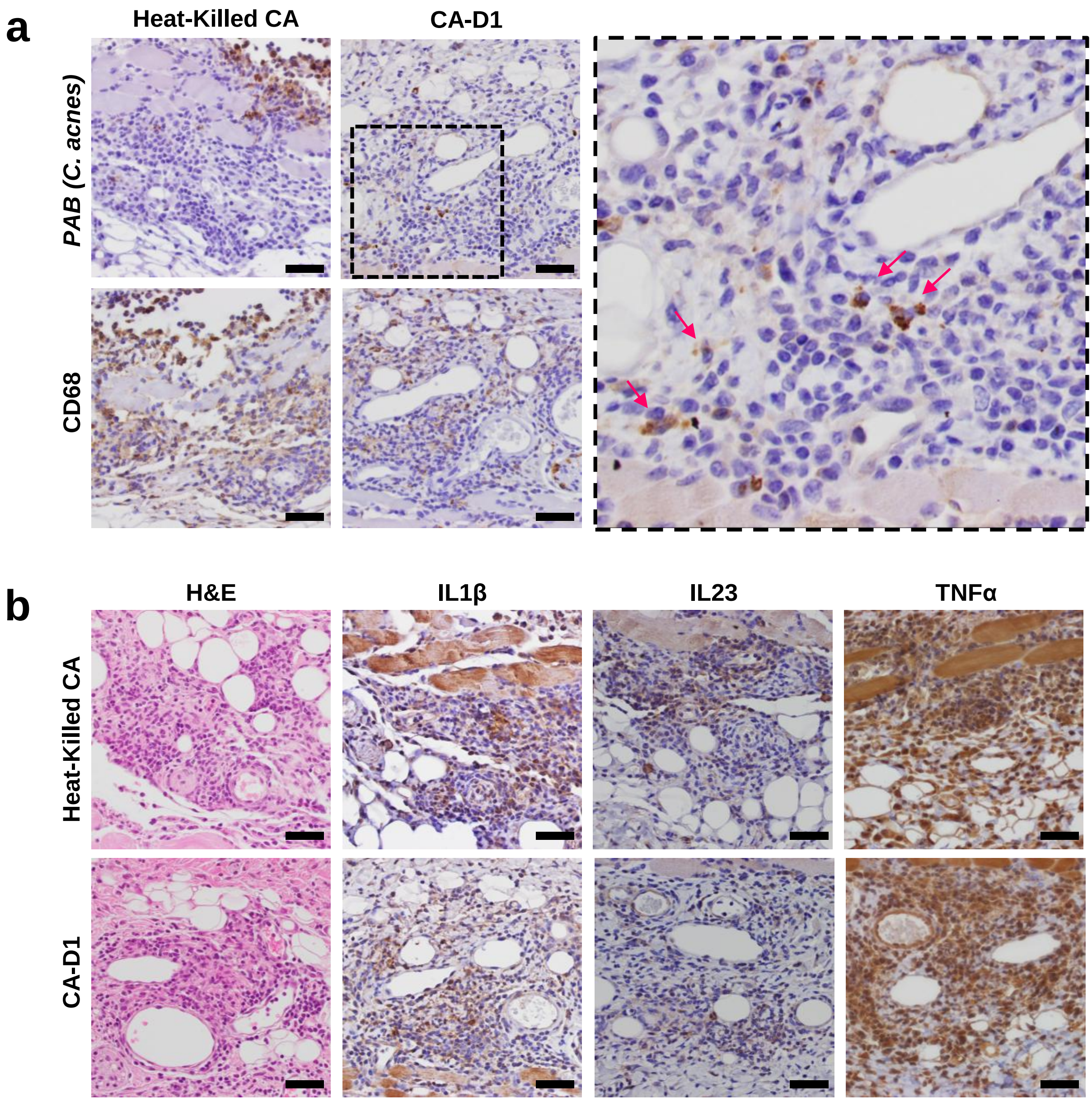


**Extended Data Fig. 6. CA-D1 and HK-CA induce aggregation of Mϕ with inflammatory responses in vivo**. **a**. IHC staining showing CA (PAB) and CD68$^{+}$ macrophages in tissues injected with HK-CA or CA-D1. Punctate CA signals were detected within Mϕ-aggregated inflammatory areas in enlarged image (right panel). (Scale bars, 20 μm) **b**. Representative H&E and IHC staining for IL-1β, IL-23, and TNF-α demonstrating similar inflammatory responses induced by HK-CA and CA-D1. (Scale bars, 20 μm)

**Extended Data Fig. 7.**

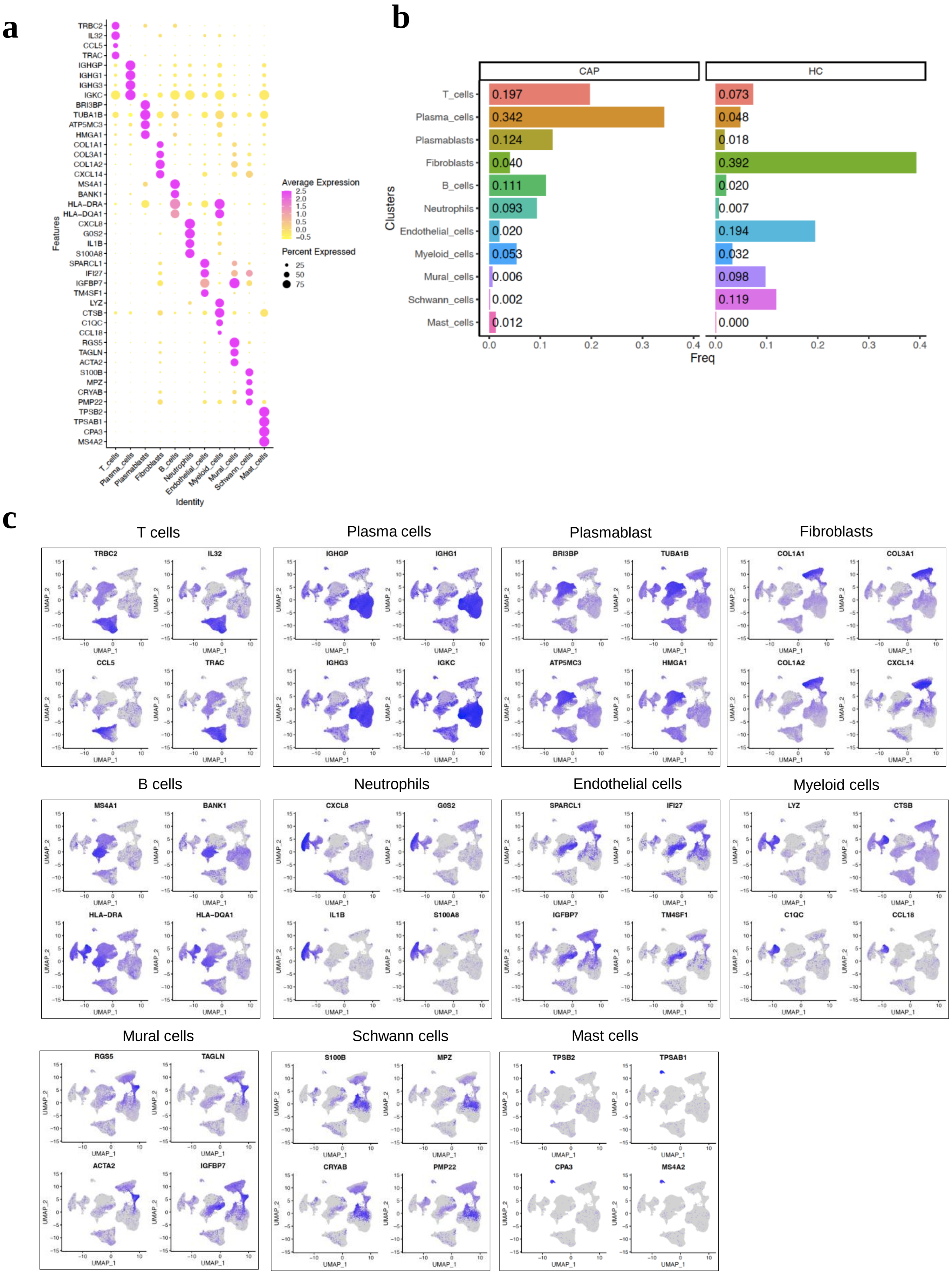

**Extended Data Fig. 7. continued**

**d**

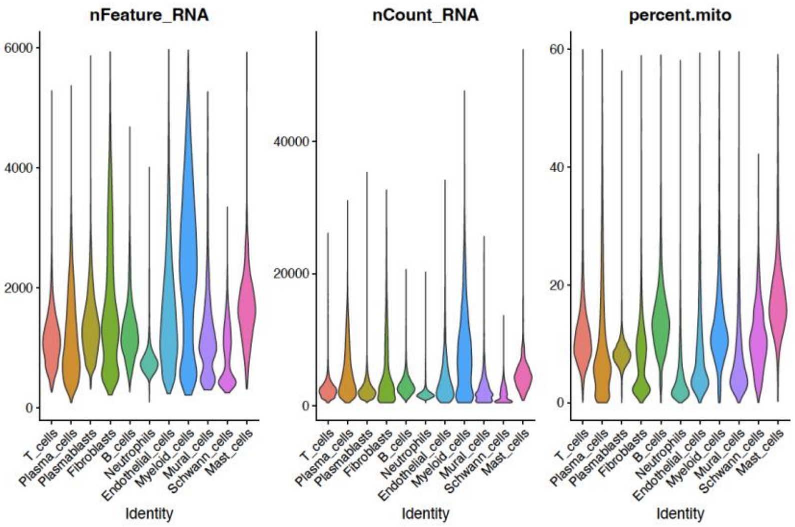


**Extended Data Fig. 7. UMAP of total cells obtained from scRNA-seq of CAP lesions and healthy control**. **a**. Dot plot showing normalized expression and the percentage of cells expressing top 4 marker genes for each cell type identified in Fig. 5b. **b**. Bar plots showing proportional cell types of CAP and healthy control (right panel). **c**. UMAP plots showing the expression of cell-type-specific marker genes from CAP and healthy control. **d**. Violin plots showing quality control metrics across major cell types, including the number of detected genes per cell (nFeature_RNA), total UMI counts per cell (nCount_RNA), and the percentage of mitochondrial gene expression (percent.mito).

**Extended Data Fig. 8.**

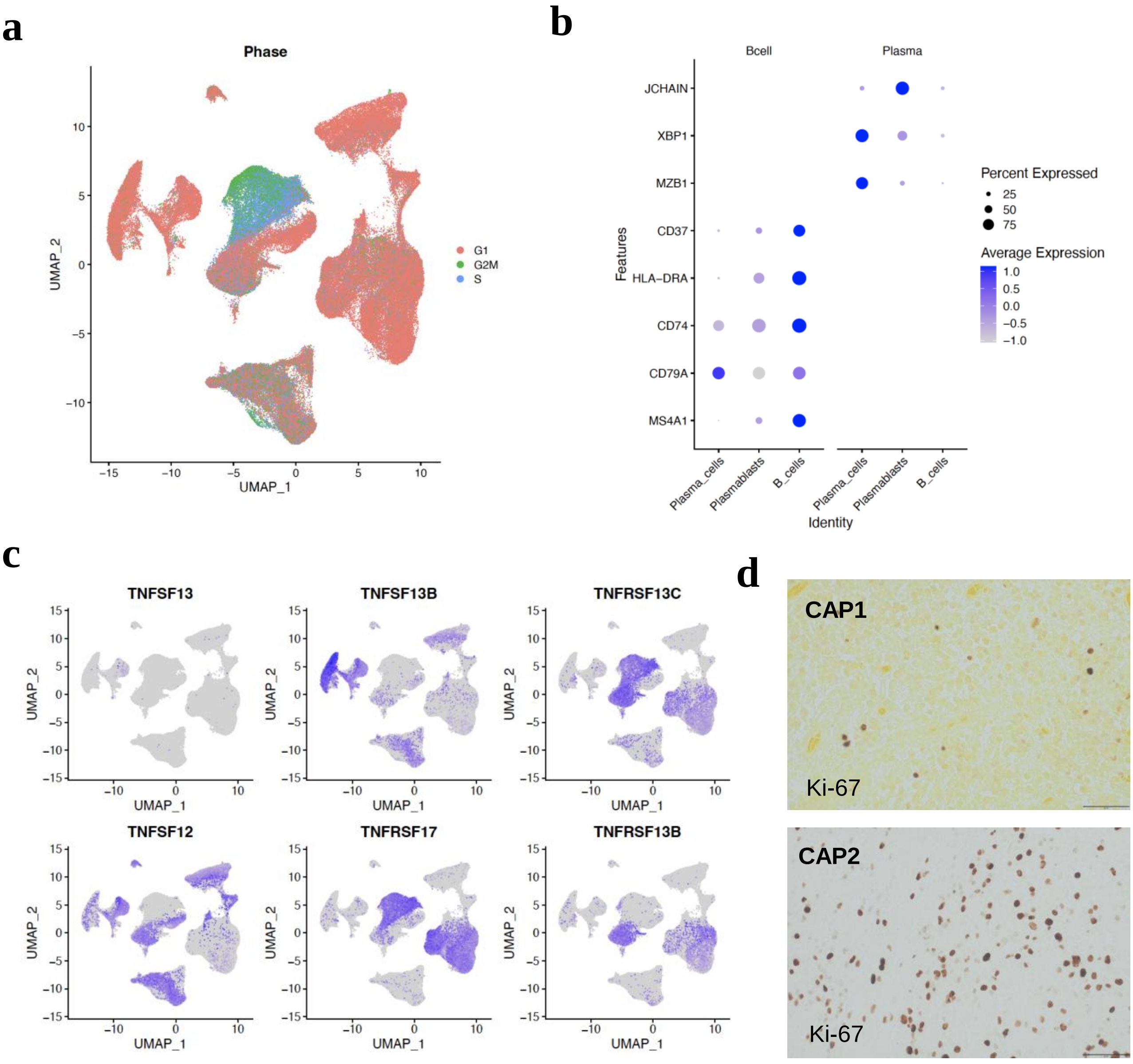


**Extended Data Figure 8. Existence of proliferating plasmablasts in CAP lesions**. **a**. UMAP plot highlighting cell cycle phase distribution in CAP lesions and healthy control tissues. **b**. Dot plot showing expression of canonical marker genes across B cells, plasmablasts, and plasma cells. Dot size represents the percentage of cells expressing each gene, and color intensity indicates average expression level. **c**. UMAP feature plots showing expression of BAFF factors in CAP lesions. **d**. Presence of proliferating immune cells in CAP lesions. Representative images of IHC staining for Ki-67 in CAP lesions.

**Extended Data Fig. 9.**

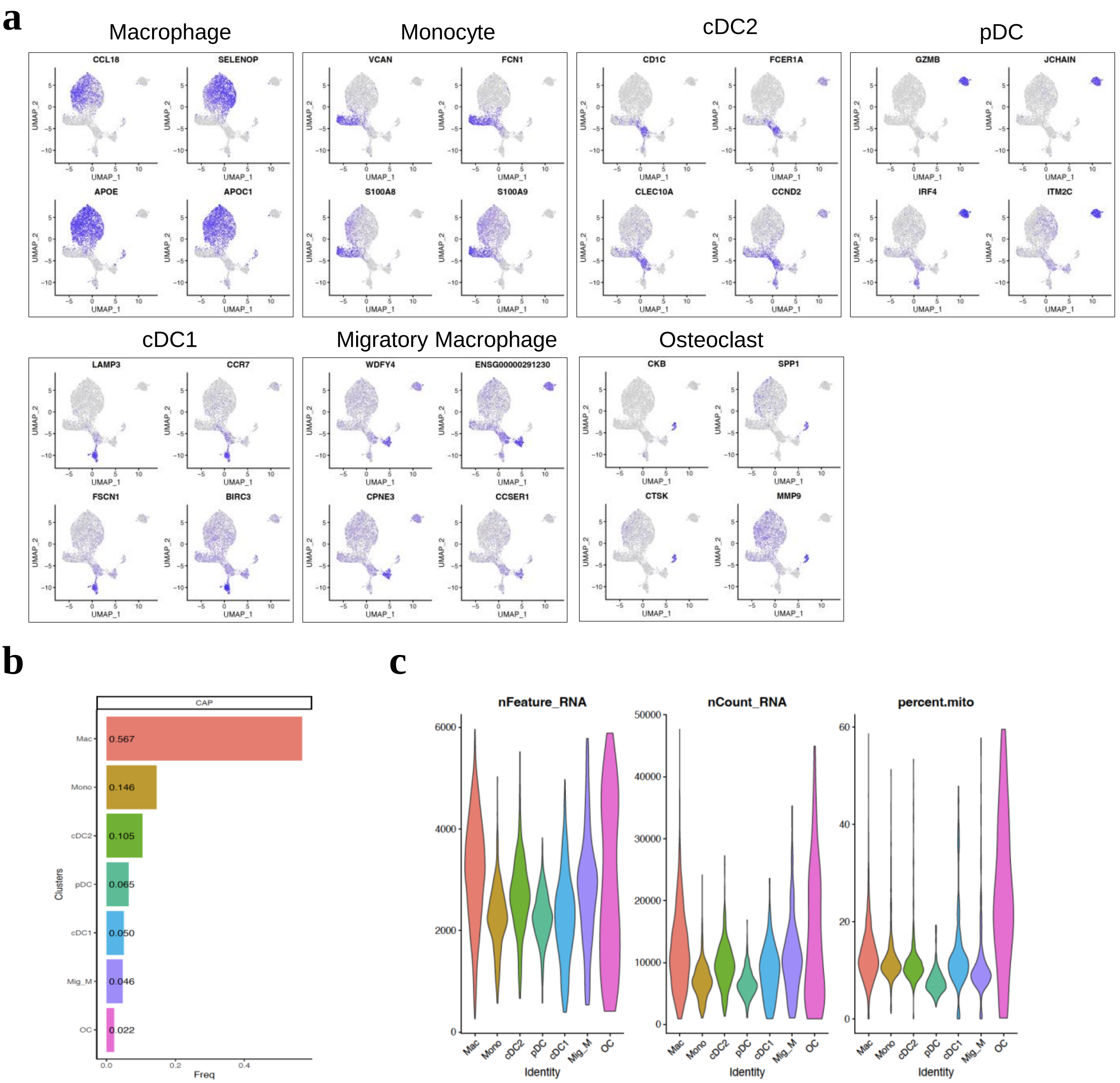


**Extended Data Figure 9. UMAP and violin plot of myeloid subsets**. **a**. UMAP feature plots showing expression of top marker genes across myeloid subsets identified in CAP lesions. **b**. Bar plot showing the relative proportions of myeloid subsets in CAP lesions. **c**. Violin plots showing quality control metrics across myeloid subsets, including the number of detected genes per cell (nFeature_RNA), total counts per cell (nCount_RNA), and percentage of mitochondrial gene expression (percent.mito).

**Extended Data Fig. 10.**

**a**

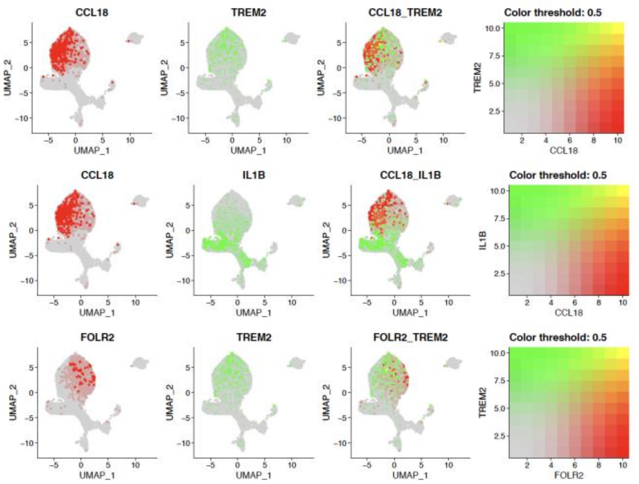


**b**

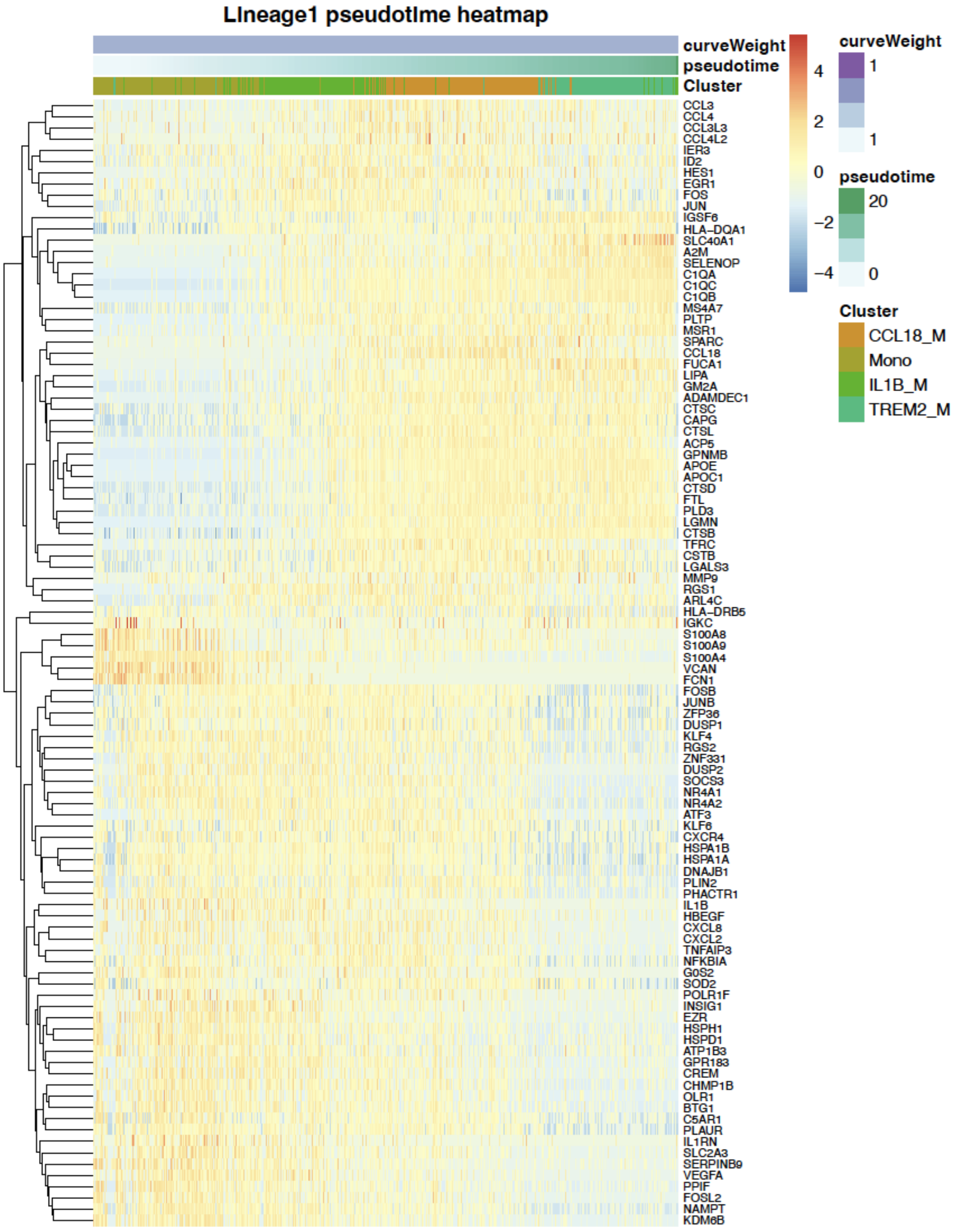

**Extended Data Fig. 10. continued**

**c**

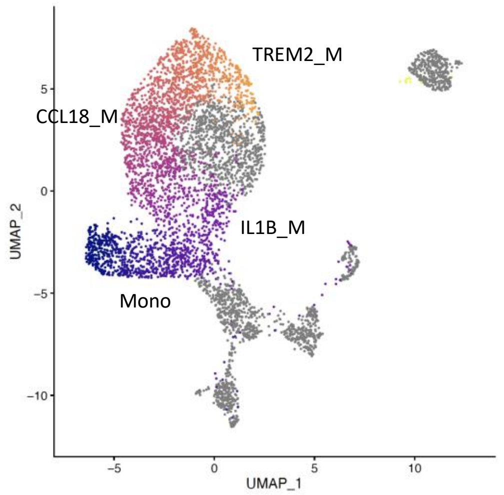


**d**

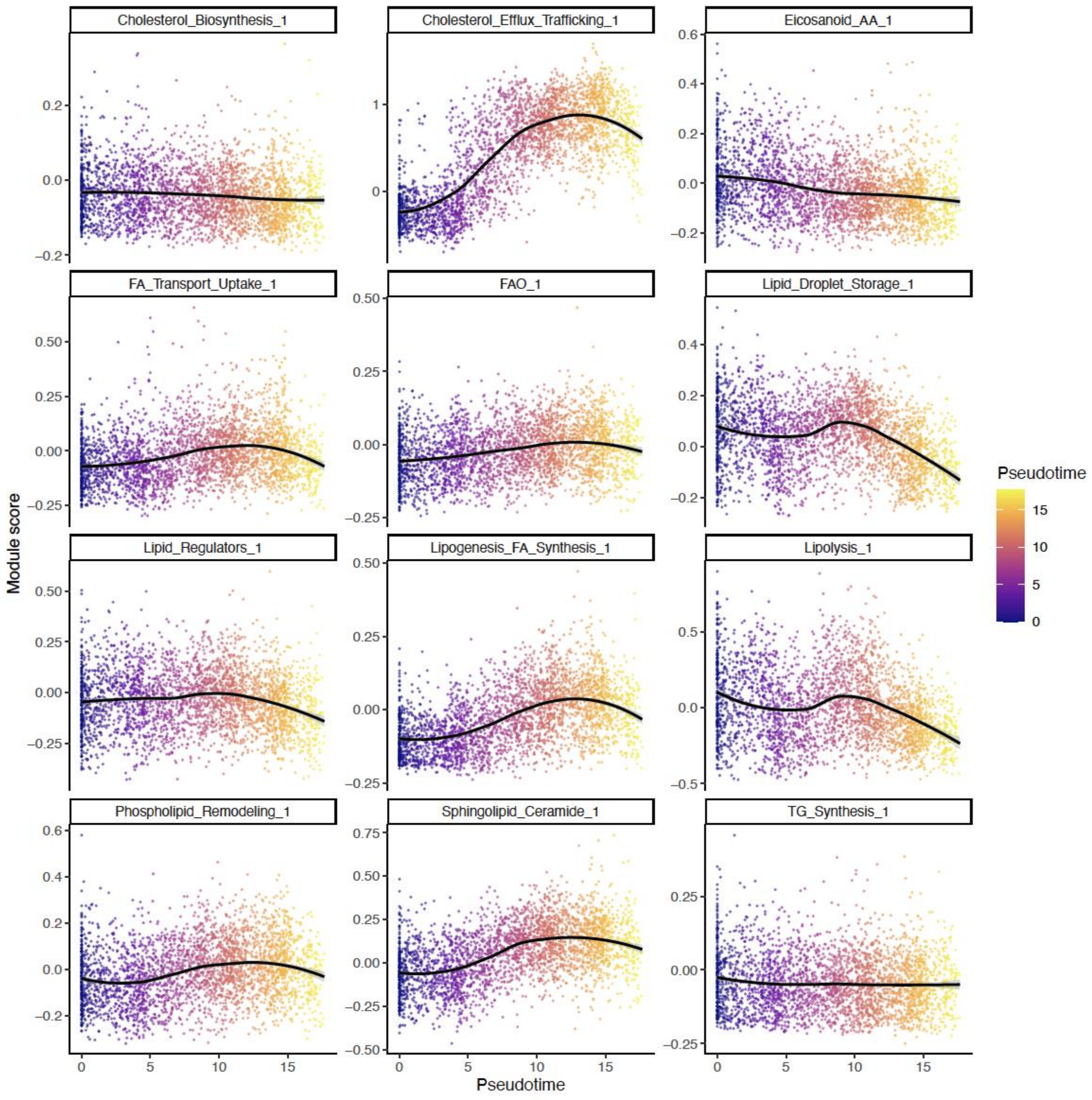

**Extended Data Fig. 10. continued**

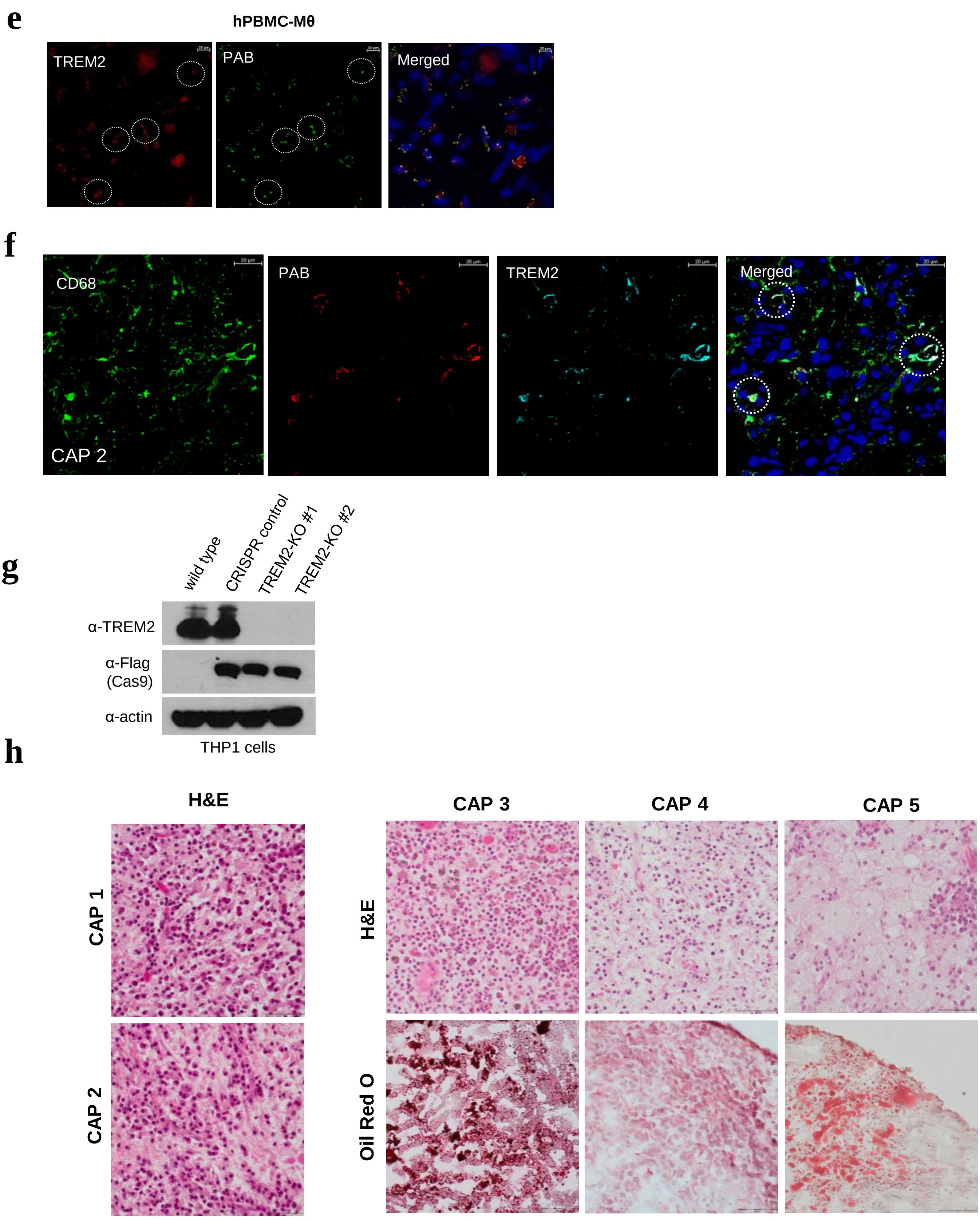

**Extended Data Figure 10. Characterization of CA persistence in TREM2 macrophages**. **a**. Feature plots showing expression patterns of CCL18, IL1β, TREM2, and FOLR2. Color scales indicate relative expression levels, and pairwise feature plots illustrate overlap between marker gene expression patterns. **b**. Heatmap showing gene expression dynamics along the inferred monocyte–macrophage trajectory in continuum. Cells are ordered based on pseudotime, with annotations indicating cluster identity. Rows represent genes, and columns represent single cells. **c**. UMAP embedding of monocyte-macrophage lineage cells colored by pseudotime, showing progression from monocytes to IL1β-high, CCL18-high, and TREM2-high macrophage subsets. **d**. Scatter plots showing gene set expression (module) scores for lipid metabolism-related pathways across pseudotime. Each point represents a single cell, colored by pseudotime, with a fitted curve indicating the overall trend. **e**. Representative IF images of CA-infected hPBMC-Mϕ for expression of TREM2 in red, CA in green. Dotted circles denote co-localization of TREM2 and CA in macrophages. (Scale bars, 10μm) **f**. Co-localization of CA and TREM2 in Mϕ detected in CAP clinical samples. CAP samples were subjected to frozen section and IF study using anti-CD68, PAB, and anti-TREM2 antibodies. Representative IF staining in CAP samples for expression of CD68 in green, CA in red, and TREM2 in cyan. (Scale bars, 20μm) **g**. Western blot analysis confirming TREM2 knockout in THP1 cells using CRISPR/Cas9-mediated gene editing. Expression of TREM2, Cas9 (Flag), and β-actin is shown. **h**. Representative images of H&E sections obtained from FFPE processing and Oil Red O staining obtained from air-dried frozen sections of CAP used in scRNA-seq analysis. H&E sections showing typical chronic inflammatory lesions with abundant plasma cells and macrophages in CAP. Note the Oil Red O staining shows a lipid-rich environment in CAP. (Scale bars, 50μm)
